# Macrogenetics reveals multifaceted influences of environmental variation on vertebrate population genetic diversity across the Americas

**DOI:** 10.1101/2021.12.15.472788

**Authors:** Elizabeth R. Lawrence, Dylan J. Fraser

## Abstract

Relative to species diversity gradients, the broad scale distribution of population-specific genetic diversity (PGD) across taxa remains understudied. We used nuclear DNA data collected from 6285 vertebrate populations across the Americas to assess the role environmental variables play in structuring the spatial/latitudinal distribution of PGD, a key component of adaptive potential in the face of environmental change. Our results provide key evidence for taxa-specific responses and that temperature variability in addition to mean temperature may be a primary driver of PGD. Additionally, we found some positive influence of precipitation, productivity, and elevation on PGD; identified trends were dependent on the metric of PGD. In contrast to the classic negative relationship between species diversity and latitude, we report either a positive or taxa-dependent relationship between PGD and latitude, depending on the metric of PGD. The inconsistent latitudinal gradient in different metrics of PGD may be due to opposing processes diminishing patterns across latitudes that operate on different timescales, as well as the flattening of large-scale genetic gradients when assessing across species versus within species. Our study highlights the nuance required to assess broad patterns in genetic diversity, and the need for developing balanced conservation strategies that ensure population, species, and community persistence.

## Introduction

Ecologists have long sought to map and understand broad patterns in biodiversity distribution, with the latitudinal gradient in species diversity being one of the most studied phenomena in ecology and biogeography. Now with increased technology and data accumulation, intraspecific diversity – such as genetic diversity within species – can be studied in the same way as species diversity. Understanding the distribution of genetic diversity is important for providing insights on the spatial distribution of biodiversity as a whole, with implications for conservation planning (De Kort et al., 2021; Lawrence & Fraser, 2020; Leigh et al., 2021; Theodoridis et al., 2020). Although still in its infancy, ‘macrogenetics’ – the study of genetic diversity across different taxa at broad scales – thus far has not revealed patterns as widespread or as obvious across latitudes as those found in species diversity (Adams & Hadly, 2013; Manel et al., 2020; Millette et al., 2020; Miraldo et al., 2016; Theodoridis et al., 2020). The environmental factors that structure spatial patterns of macrogenetics remain understudied but are likely complex and interactive (De Kort et al., 2021).

Herein we focus on the potentially non-linear and interactive effects that environmental factors have on contemporary, broad scale patterns of nuclear genetic diversity within natural populations of vertebrate species, i.e. population genetic diversity (PGD). PGD is an important component of adaptive potential in the face of changing environments. Greater genetic diversity among individuals within populations increases the likelihood of more potentially useful alleles for natural selection to act upon as the environment changes, thus increasing adaptive potential (Allendorf, 2017; Barrett & Schluter, 2008). Additionally, PGD can reliably represent geno-mewide variation through the measurement of nuclear DNA variation (Ghalambor, McKay, Carroll, & Reznick, 2007; Hurst & Jiggins, 2005; Jarne & Lagoda, 1996; Sgrò, Lowe, & Hoffmann, 2011; Väli, Einarsson, Waits, & Ellegren, 2008). Such variation is often quantifed at the population level using metrics such as heterozygosity or allelic diversity.

There are several theories that make predictions about the effect of climatic or environmental variables on PGD and the resulting spatial structuring that would occur from these effects. Some theory predicts higher PGD at low latitudes around the Equator due to their warmer temperatures and temporal climatic stability (De Kort et al., 2021; Hewitt, 2000; Schluter & Pennell, 2017). For example, higher mean temperatures are thought to increase mutation and evolution rates (Adams & Hadly, 2013; Allen, Gillooly, Savage, & Brown, 2006; Gillooly, Allen, West, & Brown, 2005; Rohde, 1992), which may favour the maintenance of higher standing PGD. Nevertheless, relationships between PGD and temperature variance over time may be more nuanced. For instance, regions that have experienced extreme temporal temperature variation can show reduced PGD due to bottlenecks (e.g. previously glaciated areas; Comps, Gömöry, Letouzey, Thiébaut, & Petit, 2001; De Kort et al., 2021; Hewitt, 2000; Schluter & Pennell, 2017). However, a great range in annual temperature may also favour higher standing PGD to deal with such unstable environments over relatively short time periods (e.g. years, decades) (Barrett & Schluter, 2008; Botero, Dor, McCain, & Safran, 2014; Brennan, Garrett, Huber, Hargarten, & Pespeni, 2019). Thus, a confounding effect between historical and contemporary ranges in temperature may exist, wherein regions that have experienced extreme temperature instability in the past may have reduced PGD overall, but regions with intermediate contemporary temperature ranges may show elevated PGD.

Other environmental factors potentially influencing broad scale PGD patterns include precipitation, productivity, and elevation. All else being equal, environments with higher precipitation and net primary productivity (e.g. at low latitudes), may support larger population sizes in many taxa. Hence such environments could maintain PGD across species because larger populations are less likely to lose genetic diversity through drift (Botero et al., 2014; Brun et al., 2019; Currie et al., 2004) (but see (Brun et al., 2019; Thuiller et al., 2020) for discussion on phylogenetic diversity and Lawrence & Fraser, 2020 for habitat area). Elevation, conversely, can be an indicator for population isolation (De Kort et al., 2021) and may interact with other environmental variables to affect PGD. For example, high elevations are typically more variable, colder environments, which can act as physiological dispersal barriers in tropical regions (Ghalambor, 2006; Janzen, 1967), and likely support smaller population sizes that experience greater genetic drift, reducing PGD. Thus, while precipitation and productivity are expected to positively influence PGD (Brun et al., 2019; De Kort et al., 2021; Santini et al., 2018), elevation is expected to have a negative and interactive effect (Ghalambor, 2006; Janzen, 1967). Together, these effects could result in higher PGD at more productive latitudes and lower elevations, such as those within 20° of the Equator (Figure 1).

**Figure 1.**
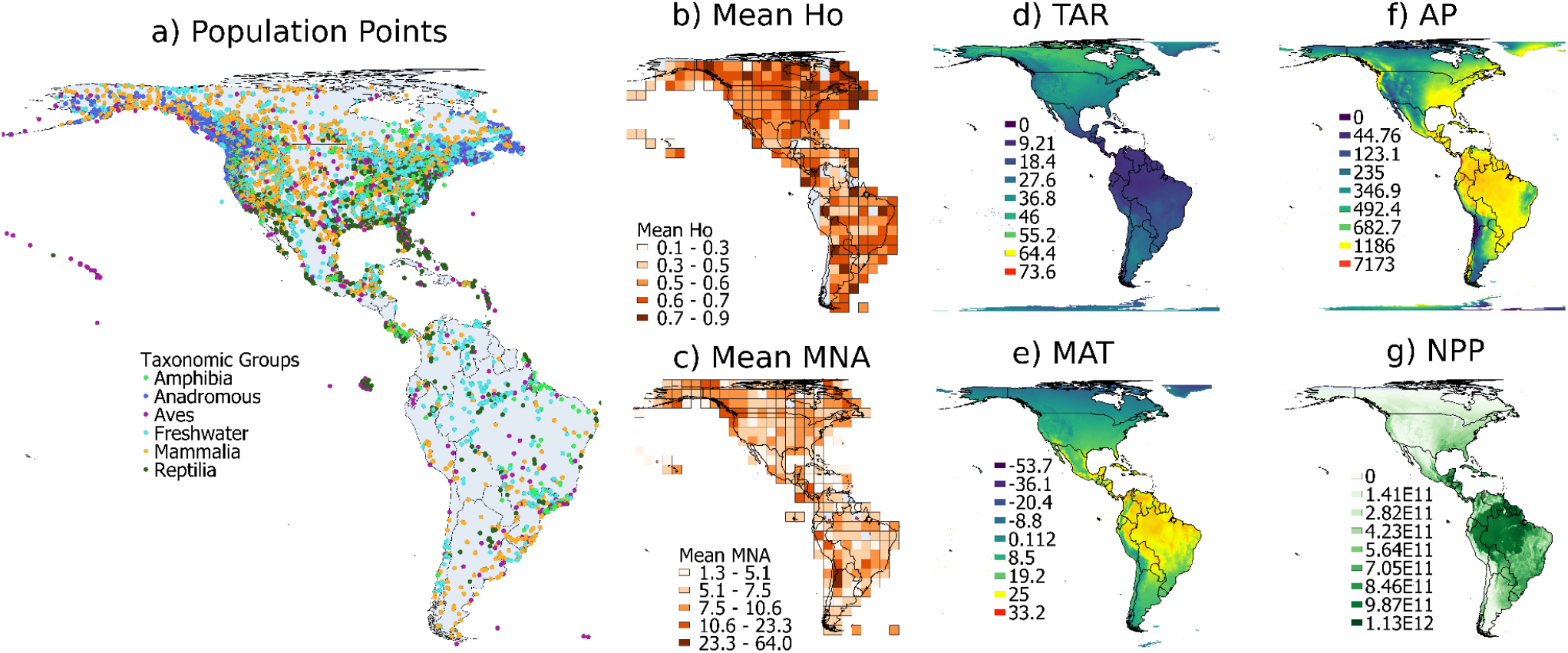
Population point localities, population genetic diversity – observed heterozygosity (Ho) and mean number of alleles (MNA) – and environmental variables across the American continents. a) Georeferenced populations coloured according to taxonomic group. Population-specific genetic diversity metrics averaged within each grid cell (b, c). Grid size is 500 km by 500 km and maps are projected in World Behrmann. d) TAR= total annual temperature range (°C); e) MAT = mean annual temperature (°C); f) AP = annual precipitation (mm/year); g) NPP = net primary productivity (units of elemental carbon x10e^-11^.

PGD has not been studied in most past macrogenetic works (Millette et al., 2020; Miraldo et al., 2016; Theodoridis et al., 2020b; Theodoridis, Rahbek, & Nogues-Bravo, 2021; but see De Kort et al. 2021). Instead, most past macrogenetic works repurposed individual-based sequence data from mitochondrial DNA. These studies found some support for a decrease in genetic diversity away from the Equator across species and taxonomic differences (e.g. stronger trends in mammals) (Adams & Hadly, 2013; Gratton et al., 2017; Martin & Mckay, 2004; Millette et al., 2020; Miraldo et al., 2016). However, inconsistencies between archiving practices in mitochondrial DNA repositories have resulted in biased measures of genetic diversity when repurposed for macrogenetic research (Leigh et al., 2021; Paz-Vinas et al., 2021). Moreover, individual-based mitochondrial DNA variation in past studies was often grouped across sampling localities in analyses (e.g. “high” or “low” latitude groups) (Adams & Hadly, 2013; Martin & Mckay, 2004; Millette et al., 2020). These limitations can generate biologically implausible data (Paz-Vinas et al., 2021) and blur critical information at the population level needed for inferring macrogenetic relationships (De Kort et al., 2021; Lawrence & Fraser, 2020). Furthermore, mitochondrial DNA variation itself has a number of shortcomings for use in macrogenetic research; for example, it may not correspond with nuclear or genome-wide genetic diversity, and it can lack resolution for demarcating populations (Allendorf, 2017; Bazin, Glemin, & Galtier, 2006; Paz-Vinas et al., 2021). Therefore, compared to nuclear DNA, mitochondrial DNA may not adequately portray broad scale genetic diversity patterns of import to conservation, such as regions harbouring populations from different species with high levels of genetic diversity for adapting to environmental change.

To determine how environmental variables influence broad scale patterns in nuclear PGD, we utilized a large database containing population-level, nuclear genetic data in vertebrate species across the American continents (Lawrence et al., 2019). This database is particularly suitable for macrogenetic research. For example, the database was generated from a systematic literature review and reports measures of putatively neutral genetic diversity for each geo-referenced, genetically distinct population (Lawrence et al., 2019). The database also includes key metadata from the original empirical studies that can be incorporated into macrogenetic analyses, such as sample sizes and multiple PGD metrics, to reduce bias and better account for nuances not considered in past macrogenetic works (Paz-Vinas et al., 2021).

Our macrogenetic study specifically used microsatellite DNA data from 6285 vertebrate populations and generalized additive mixed models to investigate environmental drivers of two metrics of PGD: observed heterozygosity (Ho) and mean number of alleles (MNA). Briefly, these metrics reflect different timescales in detecting population size reductions. Changes in MNA occur much more rapidly than in Ho, which has detectable changes over longer time scales (Allendorf, 1986; Nei, Maruyama, & Chakraborty, 1975). Therefore, our study is the first to account for possible nuances among metrics of PGD as well as non-linear relationships between PGD, latitude, and environmental factors, as raised in a recent conceptual synthesis (Lawrence & Fraser, 2020). Our study addresses two specific objectives: 1) Investigate spatial patterns of PGD by modelling Ho and MNA as a function of the following six variables: latitude, mean annual temperature (MAT), annual precipitation (AP), temperature annual range (TAR), net primary productivity (NPP), and elevation. 2) Assess taxa-specific patterns to account for differences among taxonomic groups, as different clade history and life histories may lead to differences in genetic and/or latitudinal patterns amongst groups (Adams & Hadly, 2013; De Kort et al., 2021; DeWoody & Avise, 2000; Martinez, Willoughby, & Christie, 2018; Miraldo et al., 2016).

## Methods

### Data acquisition

We extracted georeferenced vertebrate population genetic data from *MacroPopGen* that partitions microsatellite-based data for each PGD metric (Ho and MNA) (Lawrence et al., 2019). We note that the database authors already assessed and did not find a significant effect of ascertainment bias in microsatellite loci selection (Lawrence et al., 2019). We selected data from Genera that had a minimum of ten populations. For the Ho and MNA data subsets, respectively, this yielded 5143 and 4636 genetically distinct populations across the American continents from 358 and 306 species. In total, we obtained data for 6285 unique populations, 358 species, and 672 studies published between 1994 and 2017. Each population has information on Ho, MNA, sample size, and taxonomic grouping (i.e. Class, Family, Genus, Species). We obtained Ho and MNA for six vertebrate classes: anadromous and freshwater fish, amphibians, birds, mammals, and reptiles (we excluded brackish and catadromous species in *MacroPopGen* due to their low population numbers). We split fish groups into anadromous and freshwater because previous work has shown that anadromous fishes tend to show distinct genetic patterns from freshwater fishes (DeWoody & Avise, 2000; Martinez et al., 2018). We focused on Ho, instead of expected heterozygosity, because Ho is the level of genetic diversity actually observed within populations that incorporates deviations from Hardy-Weinberg expectations (Frankham, Ballou, & Briscoe, 2002). Thus, Ho offers a more realistic representation of population dynamics.

### Spatial Patterns

To test the influence of environmental variables on PGD, we extracted the following raster values based on each population data point from CHELSA (Climatologies at high resolution for the earth’s land surface areas) for the period 1979–2013: mean annual temperature (MAT, °C), annual precipitation (AP, mm/year), and temperature annual range (TAR, °C) (Karger et al., 2017a, 2017b). These contemporary environmental variables overlap roughly with the years of population-genetic data collection, so they are a reasonable estimate for the environment experienced by the populations. The elevation (m) of each geo-referenced population was obtained using the R package rgbif v1.2.0 with the srtm3 model; oceanic or aquatic regions with no data were assigned an elevation of zero. The srtm3 model is based on data collected from a sample area of 90m x 90m during the Shuttle Radar Topography Mission from the Space Shuttle Endeavour using the onboard radar system. To obtain productivity data, we extracted raster data of net primary productivity (NPP, units of elemental carbon x10e^-11^) for each population point (Imhoff & Bounoua, 2006; Imhoff et al., 2004). Finally, to identify broad spatial patterns, we mapped populations, PGD, and the rasters of environmental variables using QGIS v3.2.2 by taking the count and mean of Ho and MNA within 500km x 500km grid cells.

### Taxa-Specific Patterns

To calculate pairwise significant differences in environmental variables between vertebrate Classes (hereafter “Class”), we first performed ANOVAs (R function aov) followed by Tukey tests to correct for multiple comparisons (R function TukeyHSD) for each environmental variable. We also accounted for taxonomic influences and interactions on overall environmental patterns in models, as described below.

### Model Selection

We used generalized additive mixed models (GAMMs) to determine the effect of environmental factors on PGD using the bam() function in R package mgcv v1.8-31. We used bam over gam to improve computational performance (Pedersen, Miller, Simpson, & Ross, 2019). Response variables, Ho and MNA, were modeled separately by creating data subsets for each variable, as described above. Ho is a continuous variable bounded by zero and one and was therefore modeled with a Beta distribution; MNA values are positive, continuously distributed, and right skewed, and thus modeled with a Gamma distribution.

Genetic diversity patterns can vary among taxonomic levels (Adams & Hadly, 2013; Hirao et al., 2017). Thus, for both the Ho and MNA datasets, we included Genus as a random effect and Class as a fixed effect to interact with environmental variables. We also accounted for study as a random effect (RefID). All models were weighted by population-specific sample size of genotyped individuals to account for sample size differences among populations. Before model selection, we also tested for multicollinearity using variance inflation factors (Zuur, Ieno, Walker, Saveliev, & Smith, 2009) and the vif.gam function in R package mgcv.helper v0.1.9. No collinearity was found among variables as VIF values were less than three, below the typical threshold as indicated by (Zuur et al., 2009). In all models, continuous variables were smoothed using cubic regression splines with shrinkage applied. Shrinkage allows for a smoother to have zero degrees of freedom, thus allowing a smoother to be dropped during model selection (Zuur et al., 2009). Interaction terms were fitted with tensor products and thin-plate regression splines using the te() function in mgcv package, according to (Pedersen et al., 2019).

We used the information theoretic approach (AIC; (Akaike, 1974; Anderson & Burnham, 2002)) to conduct model selection, where the model with the lowest AIC identified the best model structure based on fit and complexity. We used forward model selection to minimize risk of overfitting models by sequentially adding one of the six environmental variables (latitude, elevation, NPP, MAT, AP, TAR) to null models until addition did not improve model fit based on AIC. We considered models within two ΔAIC points as equivalent (Zuur et al., 2009). Interactions between climatic variables may reveal micro-niche influences on PGD (Ghalambor, 2006; Janzen, 1967). For example, high elevations at low latitudes often experience similar variations in temperature variation as high latitude regions (Ghalambor, 2006; Janzen, 1967). Our model selection process therefore additionally included 11 pairwise interactions between elevation, MAT, TAR, AP, and NPP (see Table S1 for complete model structures). Finally, each environmental variable as well as latitude was interacted with Class to produce an additional six interactions during model selection. We did not include a measure of phylogenetic branch length specifically because we identified collinearity with such a variable (VIF >3), and because Class and Genus indirectly accounted for phylogenetic relationships.

Because AIC can often select overfit models (Pedersen et al., 2019; Zuur et al., 2009), we performed cross-validation on the selected models using the validation set approach with caret package v6.0-86. We assessed a variable’s significance after model selection – if a variable or interaction was not significant and/or its degrees of freedom were reduced to zero, it was removed from the model and the resulting reduced model was compared in the validation set process. We also compared several reduced models that excluded various combinations of variables (Table S3). In total, reduced models used for comparison included those that i) excluded all interactions, ii) excluded only interactions with Class, iii) excluded biologically irrelevant interactions, iv) excluded all interactions between environmental variables, v) excluded all non-significant variables and interactions, and vi) included only significant fixed effect variables. For model validation, we first trained models on a random 80% subset of the data and tested their predictive ability on the remaining 20% of the data. We calculated the Root Mean Square Error (RMSE) and the prediction error rate for selected and reduced models, selecting the model with the lowest RMSE and error rate as the final model for each PGD metric.

## Results

### Spatial Patterns

We mapped the distribution of genetically distinct vertebrate populations and their PGD, finding minimal spatial clustering in both PGD metrics (Figure 1a-c). However, grid cells with higher Ho occurred more throughout Canada and somewhat in Brazil (Figure 1b), while MNA was higher in grid cells on the western coast of North America (Figure 1c). As expected, MAT, AP, and NPP all had peak values at low latitudes centered around the equator (particularly in Brazil for NPP), whereas TAR was lowest at these latitudes (Figure 1d-g).

### Taxa-Specific Patterns

Among vertebrate classes, mean PGD and mean environmental variables varied extensively (Figure S1, Table S3, S4). Anadromous fishes showed significantly higher mean Ho (0.70) and MNA (14.97) and tended to have populations at higher latitudes (mean latitude 50.55) (p<0.001, Table S4, S5). Reptiles, birds, and amphibians experienced the highest MAT with means of 17.97, 14.25, and 12.51°C respectively. These three groups also experienced high AP, with means of 1113, 1230, and 1258 mm/year, respectively. Birds and reptiles showed the lowest TAR (22.20 and 23.19°C). Amphibians experienced the highest mean NPP (3.78e^11^ units of elemental carbon) whereas anadromous fishes experienced the lowest NPP (2.56e^11^ units of elemental carbon). Amphibians occurred at the highest mean elevation (806m), while anadromous fishes occurred at the lowest mean elevation (223m). For the complete set of pairwise comparisons and significance tests, see Table S5. Differences and interactions among Classes were also tested in the models below.

### Model Selection

The information theoretic approach (AIC) selected full models that included Class, all six environmental variables, the six interactions with Class, and 11 interactions between the environmental variables as the models of best fit for both the Ho and MNA datasets (Table S1). While all variables and interactions were significant for the Ho model (p<0.001), Class, Latitude (p=0.17), Elevation (p=0.134, NPP (p=0.25), and several interactions were insignificant for the MNA model (p>0.05; data not shown).

Beginning with the Ho dataset for cross-validation, we compared the Root Mean Square Error (RMSE) of the selected, saturated Ho model to reduced models i-iv): i) excluding all interactions, ii) excluding only interactions with Class, iii) excluding biologically irrelevant interactions, and iv) excluding all interactions between environmental variables. RMSE values were within a maximum of 0.05 units of each other, indicating the most reduced model – i.e. model i) excluding all interactions – had the lowest error prediction rate (Table S2), corresponding to (see Table S3 for summary):

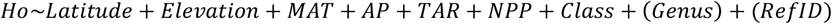

To minimize overfitting in the MNA dataset, we compared the saturated MNA model to reduced models i-vi): i) excluding all interactions, ii) excluding only interactions with Class, iii) excluding only biologically irrelevant interactions, iv) excluding all interactions between environmental variables, v) excluding all non-significant variables and interactions, and vi) including only significant fixed effects. RMSE values were within 0.01 units of each other, indicating the saturated, selected model had a slightly higher error rate than any of the reduced models (Table S2). For MNA, the model with the lowest RMSE – and thus the lowest error prediction rate – was the fourth model, corresponding to (see Table S2 for summary):

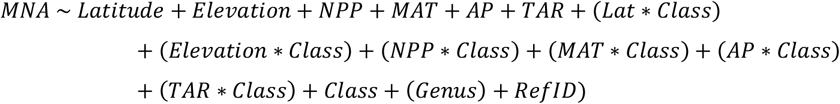

### Effect of Environmental Variables

The predicted response for PGD to each environmental variable is shown in Figure 2 and 3 for Ho and MNA, respectively; models are also summarized in Table S3. Ho increased non-linearly from mid to high latitudes (>20°) and was lower at latitudes below ~20° (Figure 2a). Ho non-linearly increased with TAR, albeit weakly, with most increases occurring above intermediate TAR of 20°C and 40°C (Figure 2b). MAT had a largely positive effect on Ho, although increases were non-linear, being highest at MAT >20°C (Figure 2c). Ho increased only somewhat with NPP (Figure 3d), slightly decreased non-linearly with AP (Figure 2e), and increased non-linearly with Elevation, particularly above 3000m (Figure 2f). Although we note there were fewer data points (n=68) at elevations above 3000m, which may be driving this increase at high elevations. Finally, while Ho varied between Classes (Figure S2), it was not significant in the model (Table S2) and Class did not interact with latitude or any of the environmental variables.

**Figure 2.**
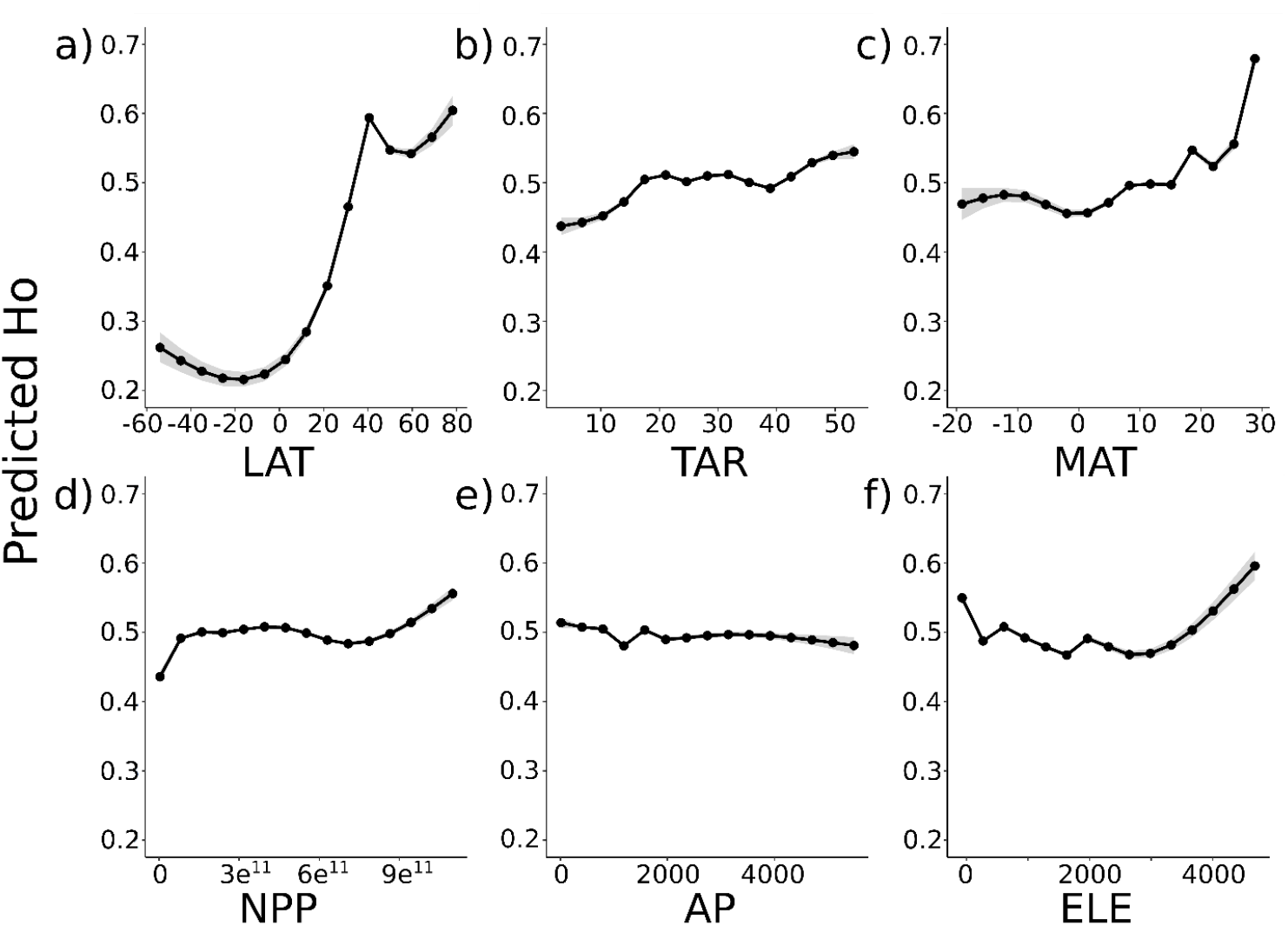
The predicted effect of latitude and environmental variables on observed heterozygosity (Ho) for species across the Americas. Predictors from the selected generalized additive mixed model were fitted by cubic smoothers and include a) degrees latitude; b), TAR= total annual temperature range (°C); c) MAT = mean annual temperature (°C); d) NPP = net primary productivity (units of elemental carbon x10e^-11^; e) AP = annual precipitation (mm/year); f) Elevation (m). Dark grey bands represent the standard error.

**Figure 3.**
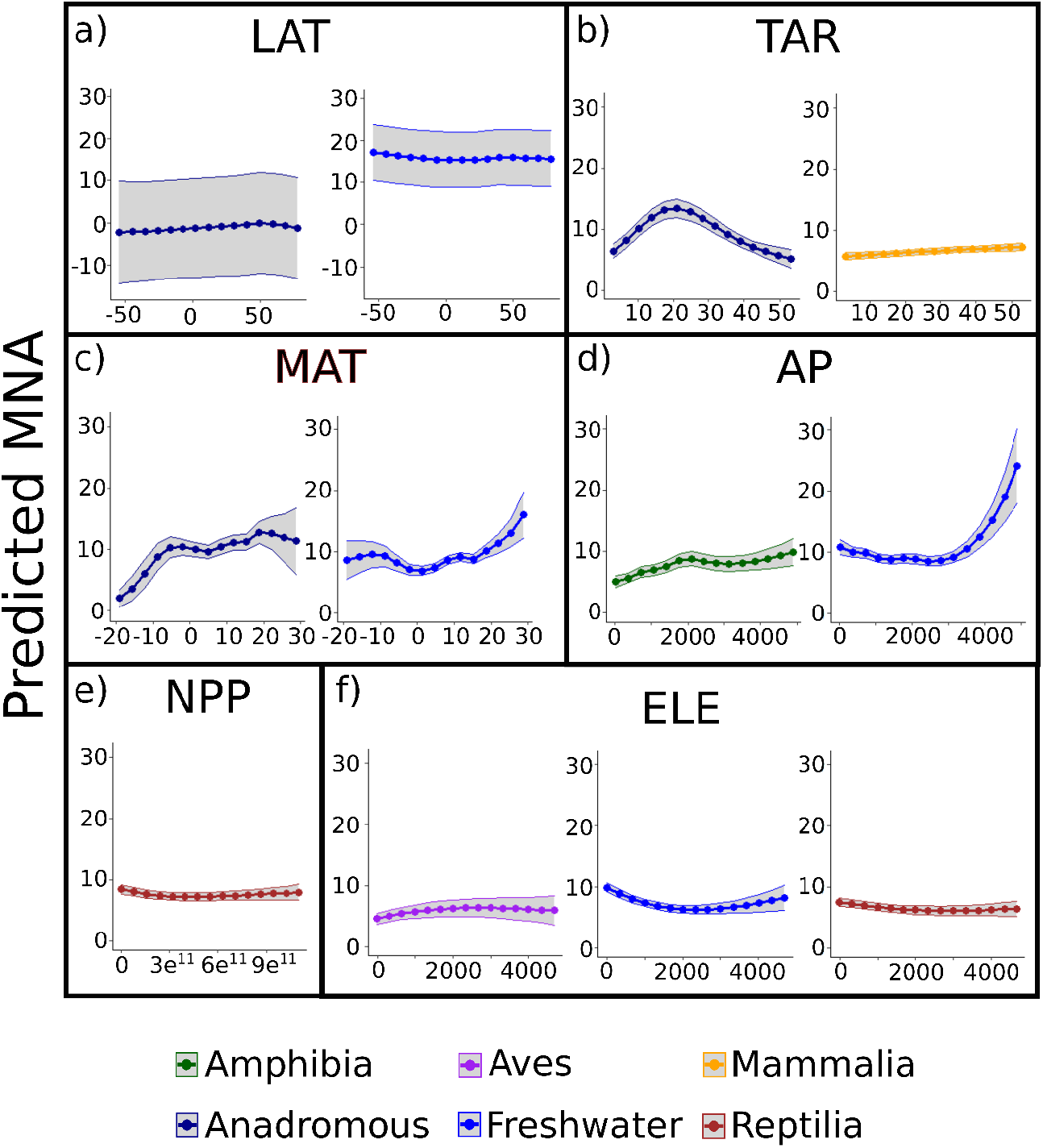
The predicted effect of latitude and environmental variables on mean number of alleles (MNA) for only the significant vertebrate Classes across the Americas (see Table S2 for model summary). Predictors were fitted by cubic smoothers and include, a) degrees latitude; b)TAR= total annual temperature range (°C); c) MAT = mean annual temperature (°C); d) AP = annual precipitation (mm/year); e) NPP = net primary productivity (units of elemental carbon x10e^-11^; f) ELE = Elevation (m). Colours represent the five different vertebrate Classes and lighter bands represent standard error. For all vertebrate groups, including non-significant relationships, see Figure S4.

Unlike Ho, the latitudinal and environmental patterns with MNA were dependent on Class (Figure 3). This was most apparent for Latitude and TAR, which showed flat relationships and did not have any significant patterns in their global smoothers (Figure S3). Even between Classes, most groups did not show any patterns with Latitude and only anadromous and freshwater fishes (Figure 3) showed any kind of relationship. MNA was vaguely U-shaped in the case of freshwater fishes, but was slightly elevated at high latitudes (<-20° and >30°) and decreased around the equator for both fish groups. Class-specific patterns varied somewhat for both TAR and MAT, where anadromous fishes differed the most, having higher MNA associated with intermediate TAR values (15 to 30°C); the remaining groups showed very little variation in MNA with TAR. MNA was highest at low MAT (−10°C) for most groups, although MNA increased above 10°C MAT for freshwater and anadromous fishes. MNA in all Classes increased slightly with NPP and AP, particularly above 3000mm/year. Finally, while MNA in amphibians, anadromous fish, and mammals did not vary with Elevation, bird MNA increased very slightly above 1000m, whereas freshwater fishes and reptile MNA decreased slightly above 1000m.

## Discussion

Our study revealed multi-faceted, taxa-dependent, and non-linear effects of environmental variables on vertebrate nuclear PGD across the American continents. These non-linear effects resulted in a lack of consistent support for a latitudinal pattern in PGD. Instead, we found that annual temperature variability and mean temperature were main variables influencing PGD. In contrast to species diversity, PGD typically increased at mid to high latitudes and decreased at low latitudes, although such patterns were not consistent between vertebrate classes. Between the two PGD metrics, we found some support for the hypotheses that MAT, TAR, AP, and NPP would have a positive effect, while Elevation would negatively affect PGD, although we again identified inconsistencies both among classes and between metrics. Interestingly, the two temperature-related variables, TAR and MAT, had non-linear effects on both metrics of PGD, with the most evident trends being increasingly higher PGD at the transition between low to intermediate TAR, and higher PGD at lower MAT (−10°C) or very high MAT (>20°C).

Temperature has been routinely suggested as a key factor that structures the latitudinal gradient in mitochondrial DNA variation, where higher temperatures increase mutation rates at the tropics, and temporal stability positively influences genetic diversity (Adams & Hadly, 2013; De Kort et al., 2021; Manel et al., 2020; Theodoridis et al., 2020). Here we had data anchored to the population level and a rich vertebrate taxonomic breadth to test such theories for nuclear genetic diversity. Only partially concordant with these expectations, and in direct contrast to results by (De Kort et al., 2021), we found that PGD was higher at lower MAT (~-10°C for MNA), the very highest MAT (>25°C for Ho and some Classes for MNA), and intermediate to high values of TAR (~20°C) (Figure 2). These results firstly indicate that warmer temperatures do not always have a positive effect on PGD – if anything, we found more evidence to the contrary. Moreover, the results are consistent with some theory that populations experiencing greater ranges in temperature, i.e. higher TAR, require higher PGD to deal with such environmental fluctuations (Barrett & Schluter, 2008; Botero et al., 2014; Brennan et al., 2019). That low MAT was usually associated with higher PGD could be due to the positive effect of higher TAR, given that MAT and TAR are negatively correlated (−0.73, Figure S5). Interestingly, for the full GAMMs that included MAT, TAR, and their interaction, the effect of the interaction was opposite for Ho and MNA: high MAT combined with low TAR positively affected MNA, but negatively affected Ho (Supplementary Files 2 and 3). This result hints at the differences between metrics of PGD and was additionally made possible by using population-specific genetic diversity, rather than aggregating potentially unrelated sequences across spatial scales.

The other environmental factors affecting PGD, were AP, NPP, and elevation, contributing to the view that the distribution of PGD is influenced by numerous factors (De Kort et al., 2021). As predicted, we found positive effects, albeit weak ones, associated with AP and NPP – although the effect of AP was only evident for MNA, particularly for freshwater fishes (Figure 3). The more evident positive relationship between AP and MNA in freshwater fishes might relate to the strong role that AP plays on freshwater habitat availability and stability (Humphries, King, & Koehn, 1999; Regier & Meisner, 1990) and/or associated adaptations required for living in such environments (Santini et al., 2018). The effect of elevation differed between the two PGD metrics, where Ho tended to increase, but MNA did not show much variation with elevation; both results contrast with our expectation that elevation should negatively influence PGD. We speculate that populations at high elevations have simply adapted to the environment and/or may not experience the typical effects associated with presumably smaller population size and greater physical isolation. Data on corresponding population size would be invaluable to test such a hypothesis but were unavailable for our dataset or for such a broad, continental scale. Overall, however, AP, NPP and Elevation did not have very strong effects on PGD relative to temperature variables.

Furthermore, a leading factor determining what affects PGD appears to be taxa-dependent, particularly for MNA. Although Ho models did not include the interactions with Class, MNA did and our results revealed that patterns may be driven by only a few Classes. For example, four of the six Classes in the MNA model showed an insignificant (i.e. flat) relationship for TAR (Figure S4). Finally, it was the two fish groups that tended to show the most significant relationships between PGD and environmental variables (7 of 12 significant relationships), suggesting these aquatic groups may respond differently from typical expectations relative to the terrestrial groups. Conversely, there may be variation within the other groups that obscure any broader-scale patterns here (e.g. mammals (Habrich, Lawrence, & Fraser, 2021)). Future works should aim to investigate such within-group variation, as we were unable to account for variability at or below the genus level due to significant loss of data.

Combined with De Kort et al. (2021), our results show that the latitudinal distribution of nuclear PGD is less clear than species diversity or previously used mitochondrial DNA data (Miraldo et al., 2016). Distinctively, our results indicate that temperature variability, along with mean temperature, might be one of the main driving factors affecting PGD. If genetic diversity is a precursor for species diversity as previous work has suggested (Adams & Hadly, 2013; Schluter & Pennell, 2017), then we would expect more PGD at latitudes and in taxa where a large degree of speciation has not yet taken place – i.e. temperate latitudes. Indeed, we found elevated Ho at latitudes >30°, and there is previous evidence for speciation rates increasing at higher latitudes (Schluter & Pennell, 2017). High latitude species also tend to experience a wider range of temperatures, or a greater TAR, and we found evidence for increasing PGD with TAR. However, there were some differences between our two PGD metrics and between vertebrate Classes. The main difference being that MNA varied much more between classes than Ho, particularly in the fish groups. Because changes to MNA occur more rapidly than in Ho, our results could reflect Class-specific differences in which vertebrates respond genetically to their environment. In a way, MNA might be a better metric for monitoring short-term PGD because it is more sensitive to changes in population size, while Ho will typically remain stable for much longer (Allendorf, 1986; Nei et al., 1975). Additionally, the differences between Ho and MNA may suggest a breakdown of latitudinal patterns in PGD, just like the changes in speciation rates. This may account for the general lack of latitudinal pattern in MNA while still finding one for Ho – insufficient time has elapsed to observe a flattening of a gradient in both metrics. We also acknowledge that anthropogenic impacts, such as urban fragmentation, may reduce PGD even more (Goossens et al., 2006) and could help explain higher Ho at latitudes above the equator (i.e. >40°), which are relatively undisturbed.

Past works identifying latitudinal gradients in genetic diversity generally did so for individual species, which may be explained by the *a priori* grouping these studies used, where species and populations were split into low or high latitude groups (Adams & Hadly, 2013; Martin & Mckay, 2004). A latitudinal gradient would appear when assessing individual species, but when looking across all species, latitudes considered low for one species may be considered high for another. Thus the latitudinal gradient is less pronounced across Classes (De Kort et al., 2021; Millette et al., 2020; Miraldo et al., 2016; Schluter & Pennell, 2017), particularly when accounting for species identity (De Kort et al., 2021; Gratton et al., 2017). This could also explain why we found little evidence for the “classic” latitudinal gradient across species.

Although our study’s dataset is taxonomically or numerically richer than past related works (Adams & Hadly, 2013; Martin & Mckay, 2004; Miraldo et al., 2016), most data (~85%) comes from North America, and there is a disproportionately large number of mammalian populations which occurred at mean latitudes of 34.19 (Table S3). This could impact interpretations of latitudinal gradients and environmental effects on PGD, as the area with the most data came from a narrower latitudinal and environmental gradient. However, when inspecting the mean PGD across North America, which is relatively well-sampled, there remained an indication of higher mean PGD at higher latitudes (i.e. western and eastern coasts of Canada for MNA and Ho, respectively; Figure 1). Additionally, there were many neutral, or flat, latitudinal relationships among Classes for the MNA model (Figure 3), suggesting PGD is not necessarily distributed latitudinally the same way species diversity is.

## Conclusion

Our results provide key evidence for taxa-dependent and different effects of environmental variables on broadscale patterns of PGD, where temperature variability and mean temperature generally had stronger, more positive effects on PGD than other environmental variables. We found positive, but weaker effects for precipitation, productivity, and elevation. We identified differences between the two genetic diversity metrics assessed – namely that MNA patterns were typically weaker than Ho and dependent on vertebrate Class, suggesting the importance of considering multiple metrics when monitoring genetic diversity. Overall, our results revealed that PGD is more structured by environmental variables than latitude alone. This result may not be surprising, but this “atypical” latitudinal gradient in PGD has important global biodiversity conservation implications. Conserving regions of high species diversity may not simultaneously conserve regions of high PGD (Paz-Vinas et al., 2018). Therefore, one must determine when the focus should remain on species richness, and when PGD should be incorporated into conservation goals to provide greater resolution. Ideally, a more comprehensive approach to conservation that considers PGD alongside species richness is recommended, as demonstrated here and in De Kort et al. (2021). We suggest that conservation goals guide the approach taken. If the goal is to conserve the greatest number of species, then species richness should be prioritized. If more narrow targets are to be met, for example the maintenance of specific species or populations, then PGD should be considered to identify which populations are either in need of prioritization or can be used to supplement other populations. Other aspects of intraspecific diversity that would be valuable to consider include population richness (Lawrence & Fraser, 2020) and adaptive genetic diversity (Lawrence & Fraser, 2020; Stanley et al., 2018). The many different populations across a species range and their genetic diversity may hold the key to future survival of the species. In sum, there is intrinsic value in managing intraspecific diversity to ensure species can adapt to and survive future environmental change (Barrett & Schluter, 2008; Bernatchez, 2016; Rey, Danchin, Mirouze, Loot, & Blanchet, 2016). Our study emphasizes this value by identifying the nuanced distribution of genetic diversity and its links to environmental factors.

## Acknowledgments

We would like to thank J-M. Matte and H. Ganz for input on the writing. Additionally, we would like to extend sincere gratitude to E. Pederson and P. Peres-Neto for their advice and guidance with the statistical approach used. Finally, we thank our funding sources which supported this work: NSERC Discovery Grant, QCBS Seed Grant, and a Concordia University Research Chair.

## Supplementary File 1

**Table S1.**
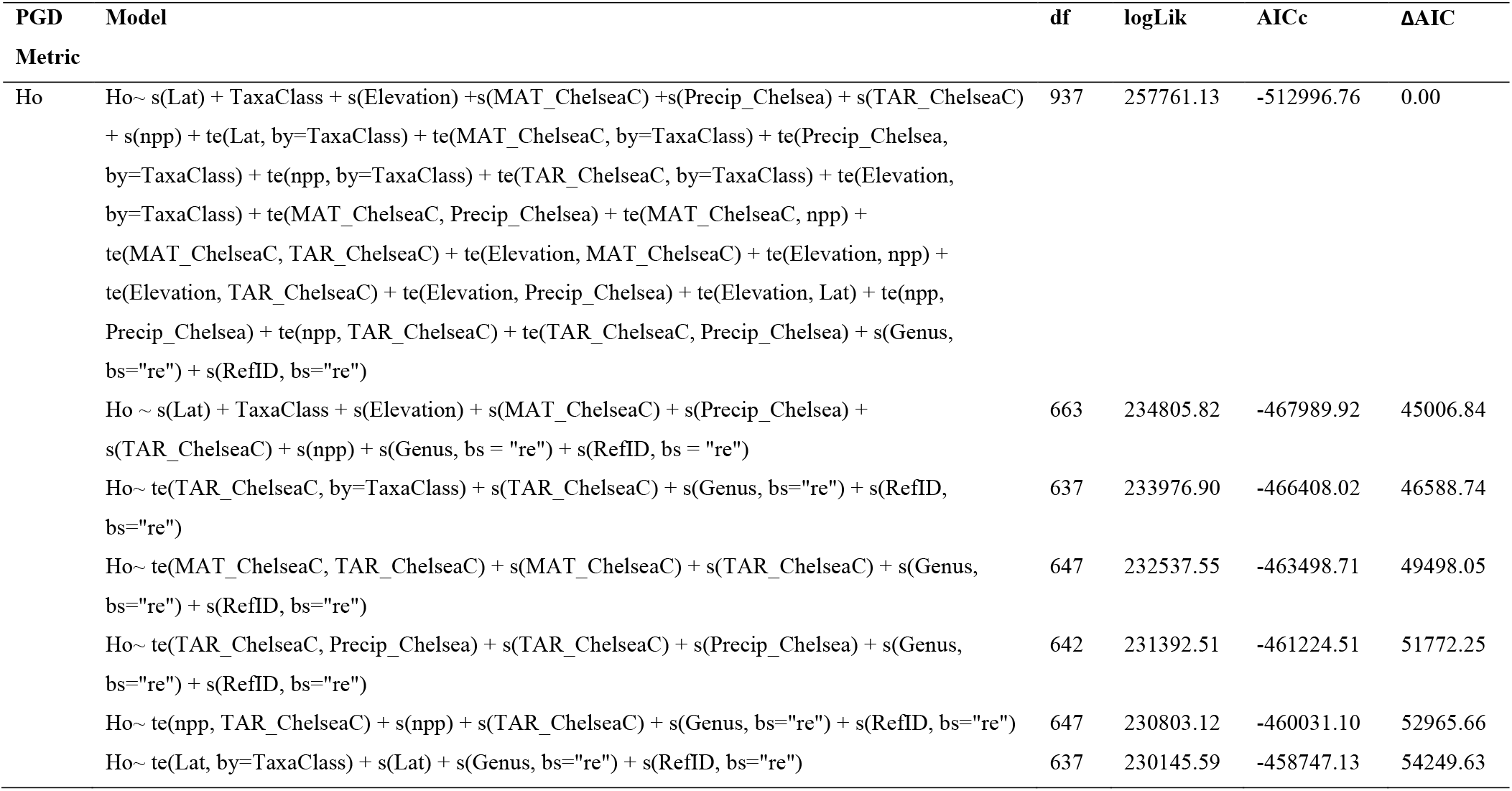
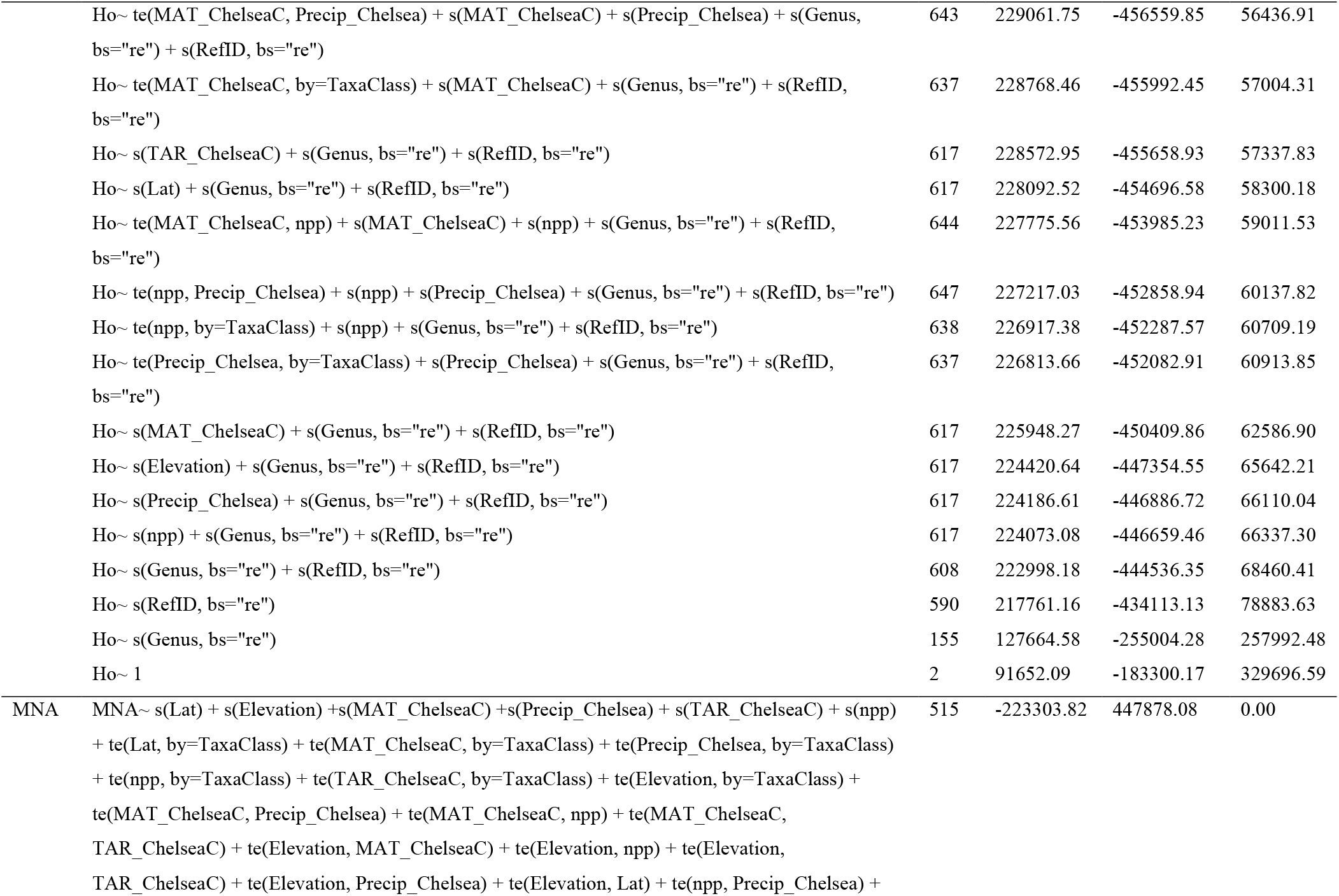
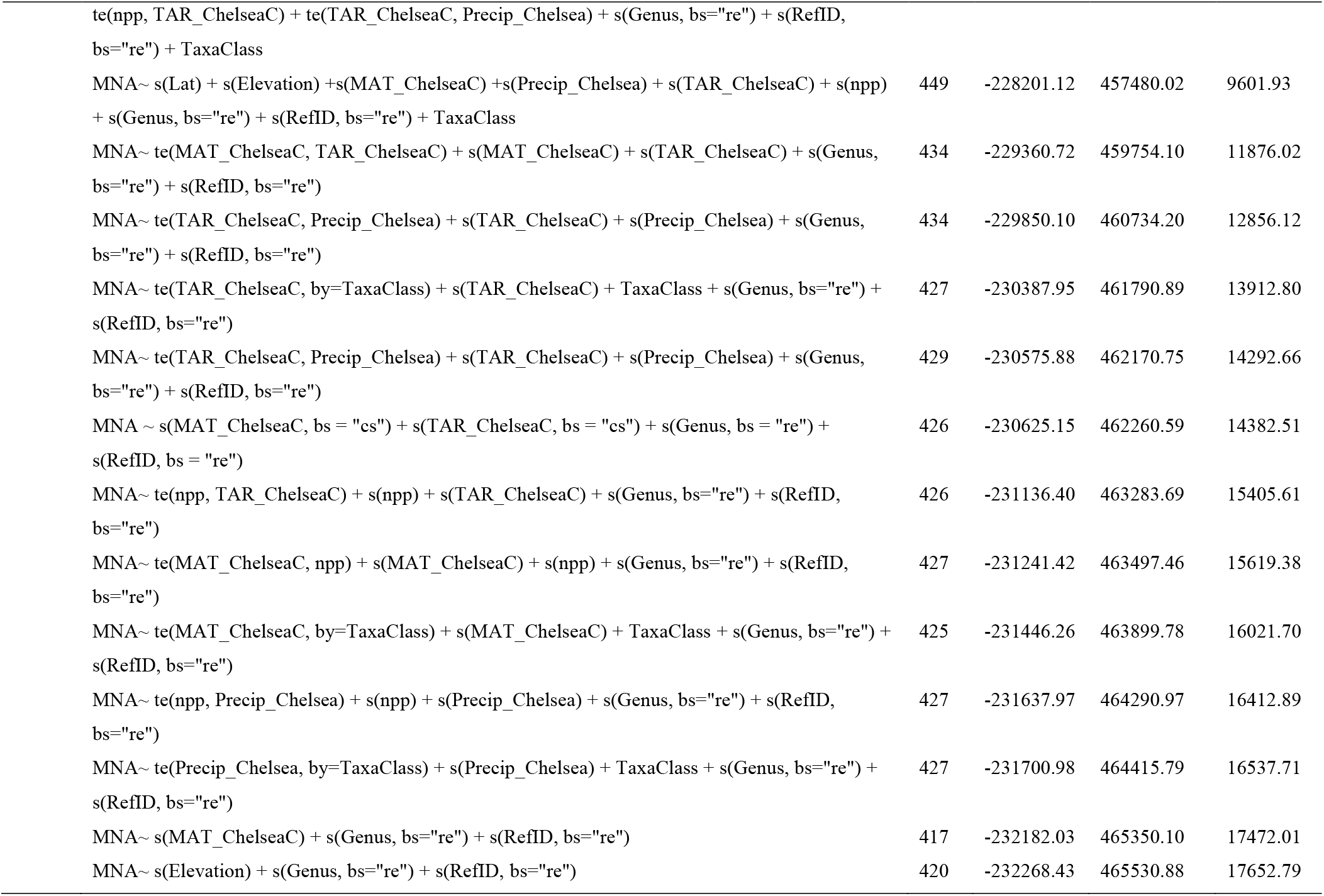
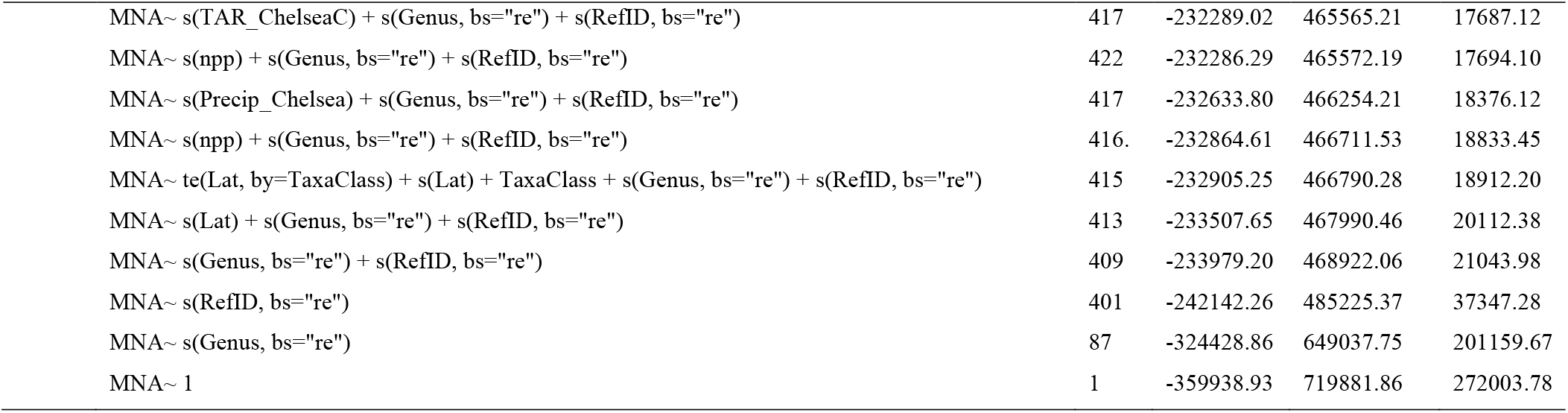
AIC comparison and model fit of select GAMMs during model selection. Ho = observed heterozygosity; MNA = mean number of alleles; Lat = degrees latitude; MAT = mean annual temperature (°C); AP = annual precipitation (mm/year); TAR = total annual range (°C); NPP = net primary productivity (units of elemental carbon x10e^-11^), Elevation (m). Response variables were fitted with smoothing parameters (s or te for interactions); random effects (bs=“re”) included Reference ID (RefID) and Genus. All models were weighted by population-specific sample size.

| PGD<br>Metric | Model | df | logLik | AICc | ΔAIC |
| --- | --- | --- | --- | --- | --- |
| Ho | Ho~ s(Lat) + TaxaClass + s(Elevation) +s(MAT_ChelseaC) +s(Precip_Chelsea) + s(TAR_ChelseaC) + s(npp) + te(Lat, by=TaxaClass) + te(MAT_ChelseaC, by=TaxaClass) + te(Precip_Chelsea, by=TaxaClass) + te(npp, by=TaxaClass) + te(TAR_ChelseaC, by=TaxaClass) + te(Elevation, by=TaxaClass) + te(MAT_ChelseaC, Precip_Chelsea) + te(MAT_ChelseaC, npp) + te(MAT_ChelseaC, TAR_ChelseaC) + te(Elevation, MAT_ChelseaC) + te(Elevation, npp) + te(Elevation, TAR_ChelseaC) + te(Elevation, Precip_Chelsea) + te(Elevation, Lat) + te(npp, Precip_Chelsea) + te(npp, TAR_ChelseaC) + te(TAR_ChelseaC, Precip_Chelsea) + s(Genus, bs="re") + s(RefID, bs="re") | 937 | 257761.13 | -512996.76 | 0.00 |
|  | Ho ~ s(Lat) + TaxaClass + s(Elevation) + s(MAT_ChelseaC) + s(Precip_Chelsea) + s(TAR_ChelseaC) + s(npp) + s(Genus, bs = "re") + s(RefID, bs = "re") | 663 | 234805.82 | -467989.92 | 45006.84 |
|  | Ho~ te(TAR_ChelseaC, by=TaxaClass) + s(TAR_ChelseaC) + s(Genus, bs="re") + s(RefID, bs="re") | 637 | 233976.90 | -466408.02 | 46588.74 |
|  | Ho~ te(MAT_ChelseaC, TAR_ChelseaC) + s(MAT_ChelseaC) + s(TAR_ChelseaC) + s(Genus, bs="re") + s(RefID, bs="re") | 647 | 232537.55 | -463498.71 | 49498.05 |
|  | Ho~ te(TAR_ChelseaC, Precip_Chelsea) + s(TAR_ChelseaC) + s(Precip_Chelsea) + s(Genus, bs="re") + s(RefID, bs="re") | 642 | 231392.51 | -461224.51 | 51772.25 |
|  | Ho~ te(npp, TAR_ChelseaC) + s(npp) + s(TAR_ChelseaC) + s(Genus, bs="re") + s(RefID, bs="re") | 647 | 230803.12 | -460031.10 | 52965.66 |
|  | Ho~ te(Lat, by=TaxaClass) + s(Lat) + s(Genus, bs="re") + s(RefID, bs="re") | 637 | 230145.59 | -458747.13 | 54249.63 |
|  | Ho~ te(MAT_ChelseaC, Precip_Chelsea) + s(MAT_ChelseaC) + s(Precip_Chelsea) + s(Genus, bs="re") + s(RefID, bs="re") | 643 | 229061.75 | -456559.85 | 56436.91 |
|  | Ho~ te(MAT_ChelseaC, by=TaxaClass) + s(MAT_ChelseaC) + s(Genus, bs="re") + s(RefID, bs="re") | 637 | 228768.46 | -455992.45 | 57004.31 |
|  | Ho~ s(TAR_ChelseaC) + s(Genus, bs="re") + s(RefID, bs="re") | 617 | 228572.95 | -455658.93 | 57337.83 |
|  | Ho~ s(Lat) + s(Genus, bs="re") + s(RefID, bs="re") | 617 | 228092.52 | -454696.58 | 58300.18 |
|  | Ho~ te(MAT_ChelseaC, npp) + s(MAT_ChelseaC) + s(npp) + s(Genus, bs="re") + s(RefID, bs="re") | 644 | 227775.56 | -453985.23 | 59011.53 |
|  | Ho~ te(npp, Precip_Chelsea) + s(npp) + s(Precip_Chelsea) + s(Genus, bs="re") + s(RefID, bs="re") | 647 | 227217.03 | -452858.94 | 60137.82 |
|  | Ho~ te(npp, by=TaxaClass) + s(npp) + s(Genus, bs="re") + s(RefID, bs="re") | 638 | 226917.38 | -452287.57 | 60709.19 |
|  | Ho~ te(Precip_Chelsea, by=TaxaClass) + s(Precip_Chelsea) + s(Genus, bs="re") + s(RefID, bs="re") | 637 | 226813.66 | -452082.91 | 60913.85 |
|  | Ho~ s(MAT_ChelseaC) + s(Genus, bs="re") + s(RefID, bs="re") | 617 | 225948.27 | -450409.86 | 62586.90 |
|  | Ho~ s(Elevation) + s(Genus, bs="re") + s(RefID, bs="re") | 617 | 224420.64 | -447354.55 | 65642.21 |
|  | Ho~ s(Precip_Chelsea) + s(Genus, bs="re") + s(RefID, bs="re") | 617 | 224186.61 | -446886.72 | 66110.04 |
|  | Ho~ s(npp) + s(Genus, bs="re") + s(RefID, bs="re") | 617 | 224073.08 | -446659.46 | 66337.30 |
|  | Ho~ s(Genus, bs="re") + s(RefID, bs="re") | 608 | 222998.18 | -444536.35 | 68460.41 |
|  | Ho~ s(RefID, bs="re") | 590 | 217761.16 | -434113.13 | 78883.63 |
|  | Ho~ s(Genus, bs="re") | 155 | 127664.58 | -255004.28 | 257992.48 |
|  | Ho~ 1 | 2 | 91652.09 | -183300.17 | 329696.59 |
| MNA | MNA~ s(Lat) + s(Elevation) + s(MAT_ChelseaC) + s(Precip_Chelsea) + s(TAR_ChelseaC) + s(npp) + te(Lat, by=TaxaClass) + te(MAT_ChelseaC, by=TaxaClass) + te(Precip_Chelsea, by=TaxaClass) + te(npp, by=TaxaClass) + te(TAR_ChelseaC, by=TaxaClass) + te(Elevation, by=TaxaClass) + te(MAT_ChelseaC, Precip_Chelsea) + te(MAT_ChelseaC, npp) + te(MAT_ChelseaC, TAR_ChelseaC) + te(Elevation, MAT_ChelseaC) + te(Elevation, npp) + te(Elevation, TAR_ChelseaC) + te(Elevation, Precip_Chelsea) + te(Elevation, Lat) + te(npp, Precip_Chelsea) + | 515 | -223303.82 | 447878.08 | 0.00 |

|  |  |  |  |  |
| --- | --- | --- | --- | --- |
| te(npp, TAR_ChelseaC) + te(TAR_ChelseaC, Precip_Chelsea) + s(Genus, bs="re") + s(RefID, bs="re") + TaxaClass |  |  |  |  |
| MNA~ s(Lat) + s(Elevation) +s(MAT_ChelseaC) +s(Precip_Chelsea) + s(TAR_ChelseaC) + s(npp) + s(Genus, bs="re") + s(RefID, bs="re") + TaxaClass | 449 | -228201.12 | 457480.02 | 9601.93 |
| MNA~ te(MAT_ChelseaC, TAR_ChelseaC) + s(MAT_ChelseaC) + s(TAR_ChelseaC) + s(Genus, bs="re") + s(RefID, bs="re") | 434 | -229360.72 | 459754.10 | 11876.02 |
| MNA~ te(TAR_ChelseaC, Precip_Chelsea) + s(TAR_ChelseaC) + s(Precip_Chelsea) + s(Genus, bs="re") + s(RefID, bs="re") | 434 | -229850.10 | 460734.20 | 12856.12 |
| MNA~ te(TAR_ChelseaC, by=TaxaClass) + s(TAR_ChelseaC) + TaxaClass + s(Genus, bs="re") + s(RefID, bs="re") | 427 | -230387.95 | 461790.89 | 13912.80 |
| MNA~ te(TAR_ChelseaC, Precip_Chelsea) + s(TAR_ChelseaC) + s(Precip_Chelsea) + s(Genus, bs="re") + s(RefID, bs="re") | 429 | -230575.88 | 462170.75 | 14292.66 |
| MNA ~ s(MAT_ChelseaC, bs = "cs") + s(TAR_ChelseaC, bs = "cs") + s(Genus, bs = "re") + s(RefID, bs = "re") | 426 | -230625.15 | 462260.59 | 14382.51 |
| MNA~ te(npp, TAR_ChelseaC) + s(npp) + s(TAR_ChelseaC) + s(Genus, bs="re") + s(RefID, bs="re") | 426 | -231136.40 | 463283.69 | 15405.61 |
| MNA~ te(MAT_ChelseaC, npp) + s(MAT_ChelseaC) + s(npp) + s(Genus, bs="re") + s(RefID, bs="re") | 427 | -231241.42 | 463497.46 | 15619.38 |
| MNA~ te(MAT_ChelseaC, by=TaxaClass) + s(MAT_ChelseaC) + TaxaClass + s(Genus, bs="re") + s(RefID, bs="re") | 425 | -231446.26 | 463899.78 | 16021.70 |
| MNA~ te(npp, Precip_Chelsea) + s(npp) + s(Precip_Chelsea) + s(Genus, bs="re") + s(RefID, bs="re") | 427 | -231637.97 | 464290.97 | 16412.89 |
| MNA~ te(Precip_Chelsea, by=TaxaClass) + s(Precip_Chelsea) + TaxaClass + s(Genus, bs="re") + s(RefID, bs="re") | 427 | -231700.98 | 464415.79 | 16537.71 |
| MNA~ s(MAT_ChelseaC) + s(Genus, bs="re") + s(RefID, bs="re") | 417 | -232182.03 | 465350.10 | 17472.01 |
| MNA~ s(Elevation) + s(Genus, bs="re") + s(RefID, bs="re") | 420 | -232268.43 | 465530.88 | 17652.79 |

|  |  |  |  |  |
| --- | --- | --- | --- | --- |
| MNA~ s(TAR_ChelseaC) + s(Genus, bs="re") + s(RefID, bs="re") | 417 | -232289.02 | 465565.21 | 17687.12 |
| MNA~ s(npp) + s(Genus, bs="re") + s(RefID, bs="re") | 422 | -232286.29 | 465572.19 | 17694.10 |
| MNA~ s(Precip_Chelsea) + s(Genus, bs="re") + s(RefID, bs="re") | 417 | -232633.80 | 466254.21 | 18376.12 |
| MNA~ s(npp) + s(Genus, bs="re") + s(RefID, bs="re") | 416. | -232864.61 | 466711.53 | 18833.45 |
| MNA~ te(Lat, by=TaxaClass) + s(Lat) + TaxaClass + s(Genus, bs="re") + s(RefID, bs="re") | 415 | -232905.25 | 466790.28 | 18912.20 |
| MNA~ s(Lat) + s(Genus, bs="re") + s(RefID, bs="re") | 413 | -233507.65 | 467990.46 | 20112.38 |
| MNA~ s(Genus, bs="re") + s(RefID, bs="re") | 409 | -233979.20 | 468922.06 | 21043.98 |
| MNA~ s(RefID, bs="re") | 401 | -242142.26 | 485225.37 | 37347.28 |
| MNA~ s(Genus, bs="re") | 87 | -324428.86 | 649037.75 | 201159.67 |
| MNA~ 1 | 1 | -359938.93 | 719881.86 | 272003.78 |

**Table S2.**
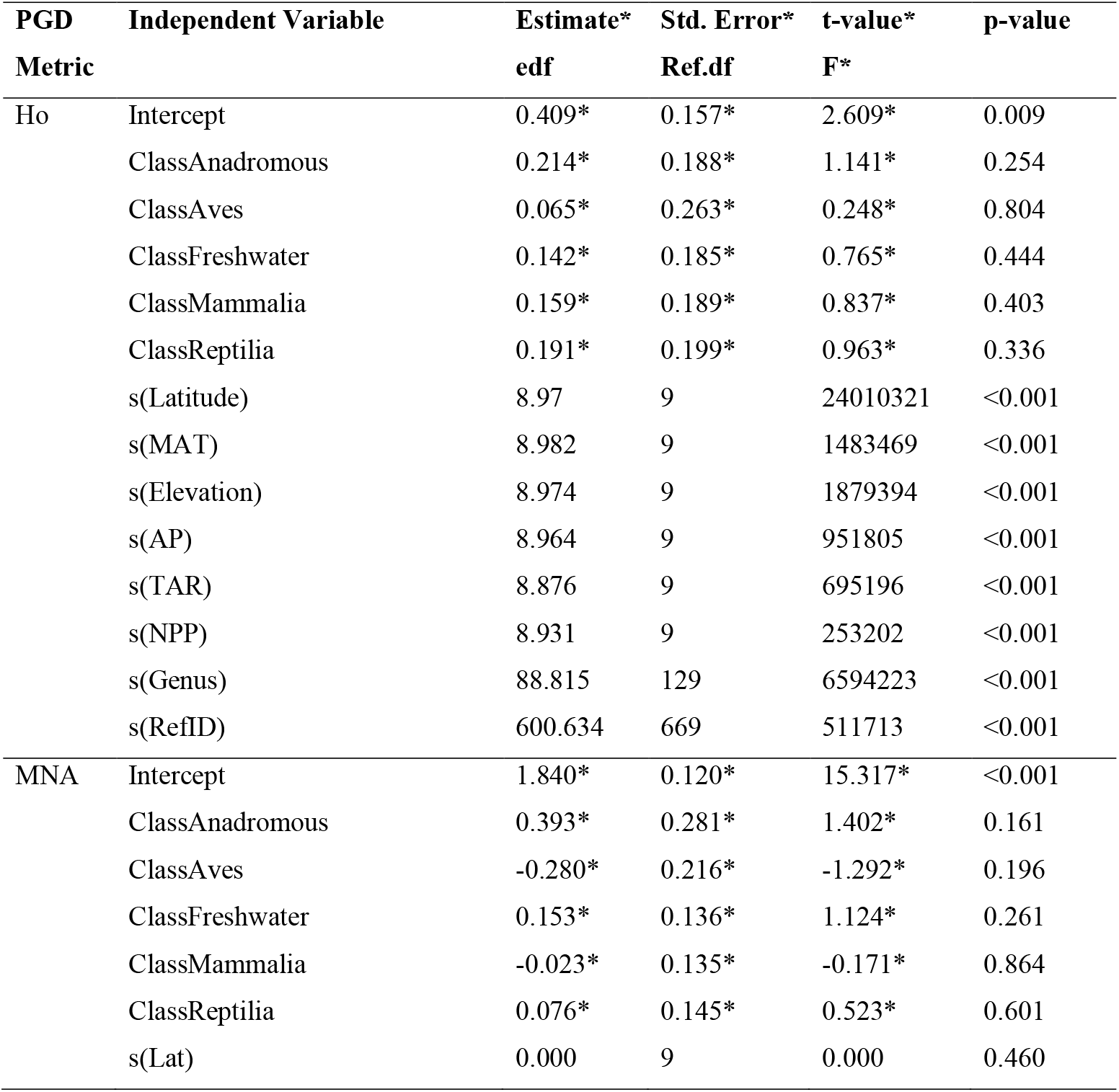
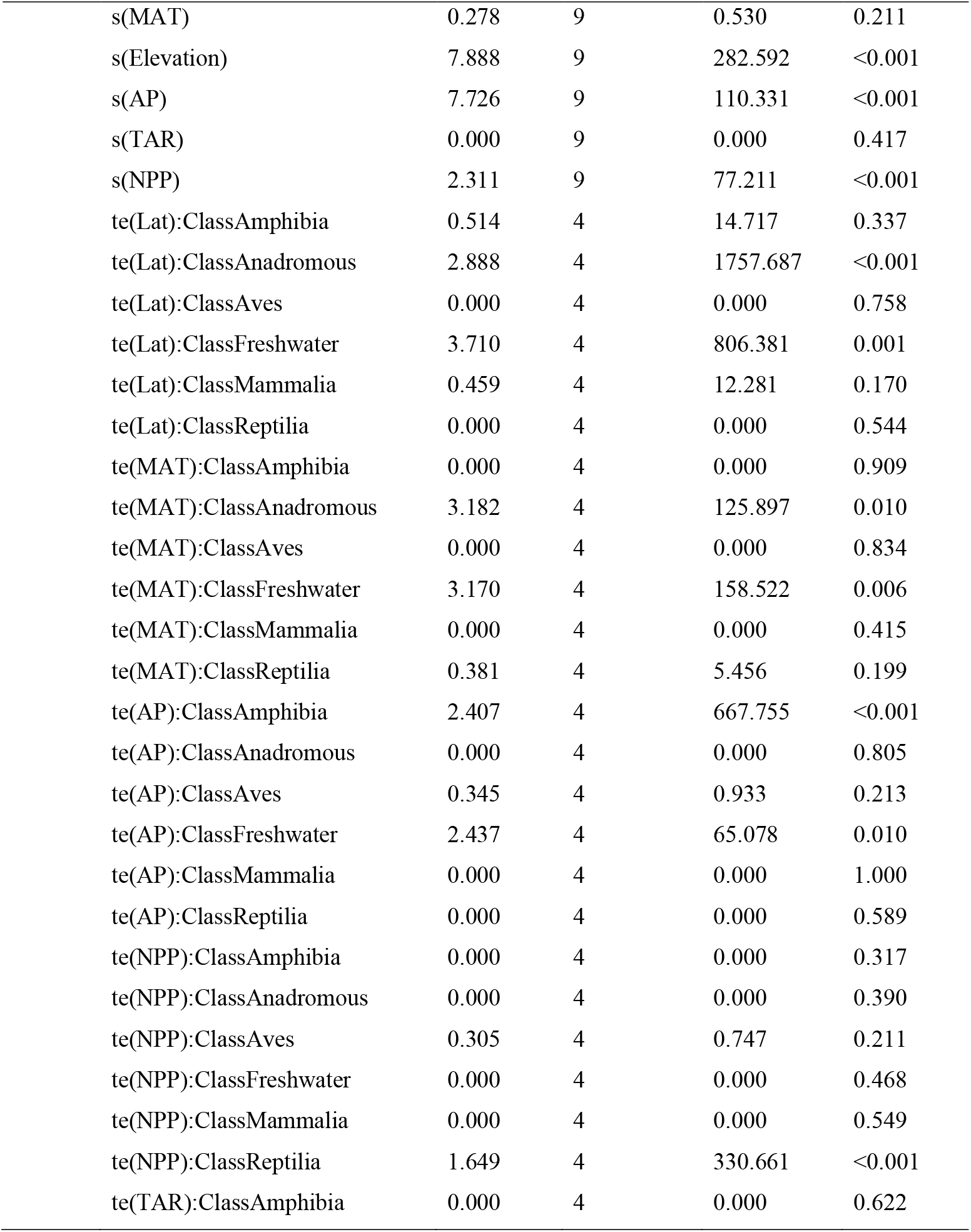
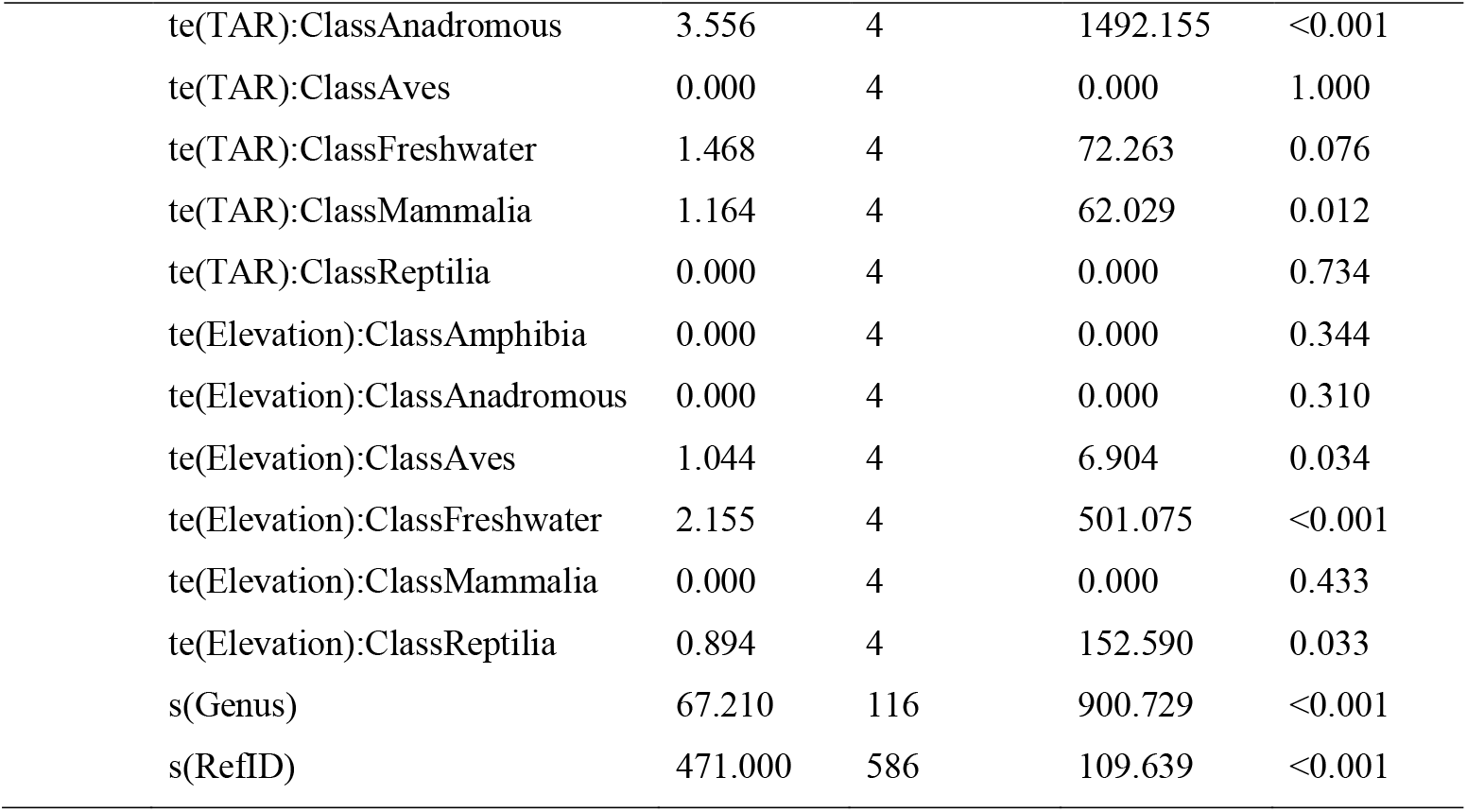
Summary of selected GAMMs for either Ho (modeled with beta distribution) or MNA (gamma distribution). Predictor variables fitted with a smoother (s) included Lat = latitude, MAT = mean annual temperature (°C), Elevation (m), AP = annual precipitation (mm/year), TAR = total annual range (°C), NPP = net primary productivity (units of elemental carbon x10e^-11^). Interactions are fitted with tensor products (te). Random effects included Genus and Reference ID (RefID).

| **PGD** | **Independent Variable** | **Estimate\*** | **Std. Error\*** | **t-value\*** | **p-value** |
| --- | --- | --- | --- | --- | --- |
| **Metric** |  | **edf** | **Ref.df** | **F\*** |  |
| Ho | Intercept | 0.409* | 0.157* | 2.609* | 0.009 |
|  | ClassAnadromous | 0.214* | 0.188* | 1.141* | 0.254 |
|  | ClassAves | 0.065* | 0.263* | 0.248* | 0.804 |
|  | ClassFreshwater | 0.142* | 0.185* | 0.765* | 0.444 |
|  | ClassMammalia | 0.159* | 0.189* | 0.837* | 0.403 |
|  | ClassReptilia | 0.191* | 0.199* | 0.963* | 0.336 |
|  | s(Latitude) | 8.97 | 9 | 24010321 | <0.001 |
|  | s(MAT) | 8.982 | 9 | 1483469 | <0.001 |
|  | s(Elevation) | 8.974 | 9 | 1879394 | <0.001 |
|  | s(AP) | 8.964 | 9 | 951805 | <0.001 |
|  | s(TAR) | 8.876 | 9 | 695196 | <0.001 |
|  | s(NPP) | 8.931 | 9 | 253202 | <0.001 |
|  | s(Genus) | 88.815 | 129 | 6594223 | <0.001 |
|  | s(RefID) | 600.634 | 669 | 511713 | <0.001 |
| MNA | Intercept | 1.840* | 0.120* | 15.317* | <0.001 |
|  | ClassAnadromous | 0.393* | 0.281* | 1.402* | 0.161 |
|  | ClassAves | -0.280* | 0.216* | -1.292* | 0.196 |
|  | ClassFreshwater | 0.153* | 0.136* | 1.124* | 0.261 |
|  | ClassMammalia | -0.023* | 0.135* | -0.171* | 0.864 |
|  | ClassReptilia | 0.076* | 0.145* | 0.523* | 0.601 |
|  | s(Lat) | 0.000 | 9 | 0.000 | 0.460 |

|  |  |  |  |  |
| --- | --- | --- | --- | --- |
| s(MAT) | 0.278 | 9 | 0.530 | 0.211 |
| s(Elevation) | 7.888 | 9 | 282.592 | <0.001 |
| s(AP) | 7.726 | 9 | 110.331 | <0.001 |
| s(TAR) | 0.000 | 9 | 0.000 | 0.417 |
| s(NPP) | 2.311 | 9 | 77.211 | <0.001 |
| te(Lat):ClassAmphibia | 0.514 | 4 | 14.717 | 0.337 |
| te(Lat):ClassAnadromous | 2.888 | 4 | 1757.687 | <0.001 |
| te(Lat):ClassAves | 0.000 | 4 | 0.000 | 0.758 |
| te(Lat):ClassFreshwater | 3.710 | 4 | 806.381 | 0.001 |
| te(Lat):ClassMammalia | 0.459 | 4 | 12.281 | 0.170 |
| te(Lat):ClassReptilia | 0.000 | 4 | 0.000 | 0.544 |
| te(MAT):ClassAmphibia | 0.000 | 4 | 0.000 | 0.909 |
| te(MAT):ClassAnadromous | 3.182 | 4 | 125.897 | 0.010 |
| te(MAT):ClassAves | 0.000 | 4 | 0.000 | 0.834 |
| te(MAT):ClassFreshwater | 3.170 | 4 | 158.522 | 0.006 |
| te(MAT):ClassMammalia | 0.000 | 4 | 0.000 | 0.415 |
| te(MAT):ClassReptilia | 0.381 | 4 | 5.456 | 0.199 |
| te(AP):ClassAmphibia | 2.407 | 4 | 667.755 | <0.001 |
| te(AP):ClassAnadromous | 0.000 | 4 | 0.000 | 0.805 |
| te(AP):ClassAves | 0.345 | 4 | 0.933 | 0.213 |
| te(AP):ClassFreshwater | 2.437 | 4 | 65.078 | 0.010 |
| te(AP):ClassMammalia | 0.000 | 4 | 0.000 | 1.000 |
| te(AP):ClassReptilia | 0.000 | 4 | 0.000 | 0.589 |
| te(NPP):ClassAmphibia | 0.000 | 4 | 0.000 | 0.317 |
| te(NPP):ClassAnadromous | 0.000 | 4 | 0.000 | 0.390 |
| te(NPP):ClassAves | 0.305 | 4 | 0.747 | 0.211 |
| te(NPP):ClassFreshwater | 0.000 | 4 | 0.000 | 0.468 |
| te(NPP):ClassMammalia | 0.000 | 4 | 0.000 | 0.549 |
| te(NPP):ClassReptilia | 1.649 | 4 | 330.661 | <0.001 |
| te(TAR):ClassAmphibia | 0.000 | 4 | 0.000 | 0.622 |

|  |  |  |  |  |
| --- | --- | --- | --- | --- |
| te(TAR):ClassAnadromous | 3.556 | 4 | 1492.155 | <0.001 |
| te(TAR):ClassAves | 0.000 | 4 | 0.000 | 1.000 |
| te(TAR):ClassFreshwater | 1.468 | 4 | 72.263 | 0.076 |
| te(TAR):ClassMammalia | 1.164 | 4 | 62.029 | 0.012 |
| te(TAR):ClassReptilia | 0.000 | 4 | 0.000 | 0.734 |
| te(Elevation):ClassAmphibia | 0.000 | 4 | 0.000 | 0.344 |
| te(Elevation):ClassAnadromous | 0.000 | 4 | 0.000 | 0.310 |
| te(Elevation):ClassAves | 1.044 | 4 | 6.904 | 0.034 |
| te(Elevation):ClassFreshwater | 2.155 | 4 | 501.075 | <0.001 |
| te(Elevation):ClassMammalia | 0.000 | 4 | 0.000 | 0.433 |
| te(Elevation):ClassReptilia | 0.894 | 4 | 152.590 | 0.033 |
| s(Genus) | 67.210 | 116 | 900.729 | <0.001 |
| s(RefID) | 471.000 | 586 | 109.639 | <0.001 |

**Table S3.**
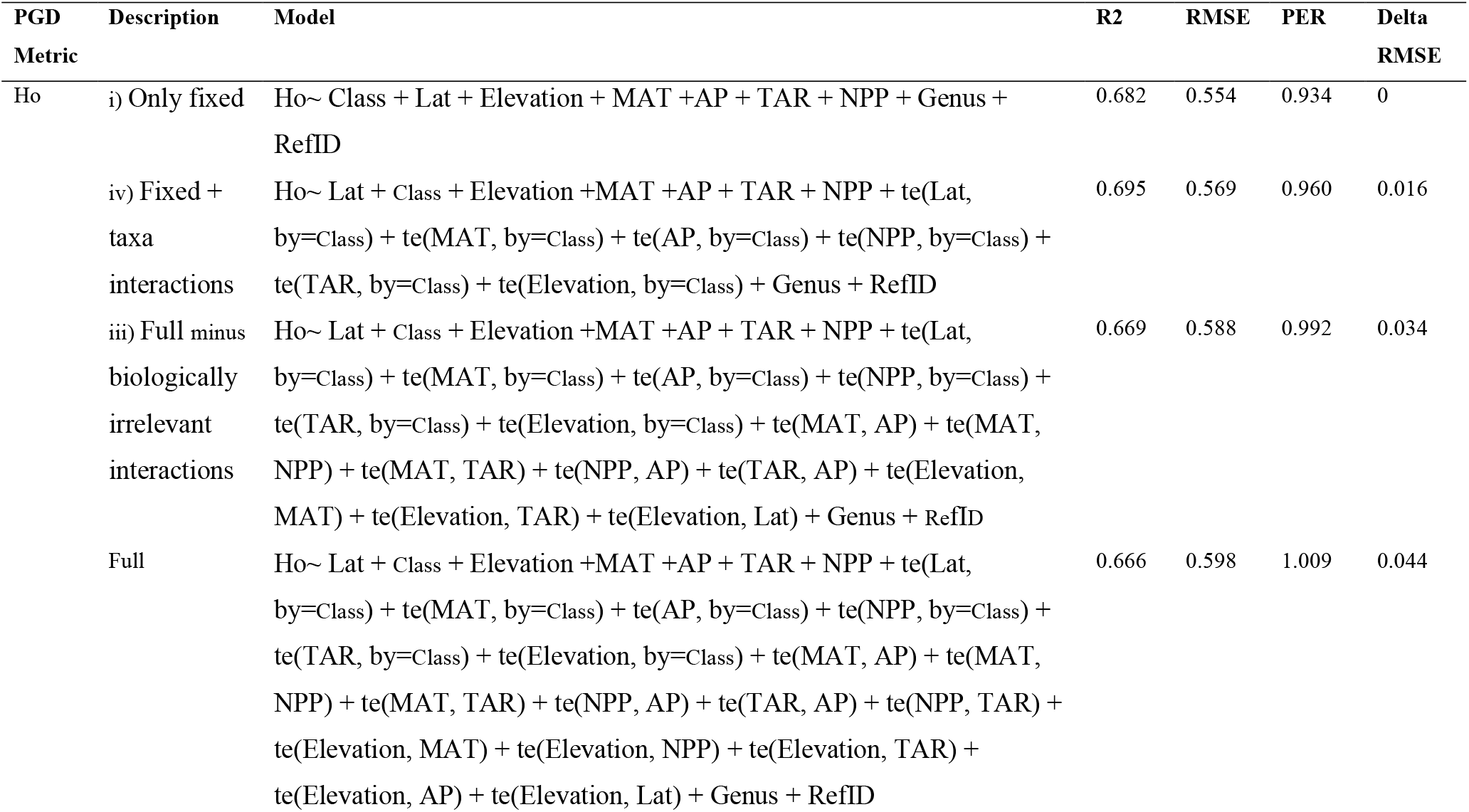
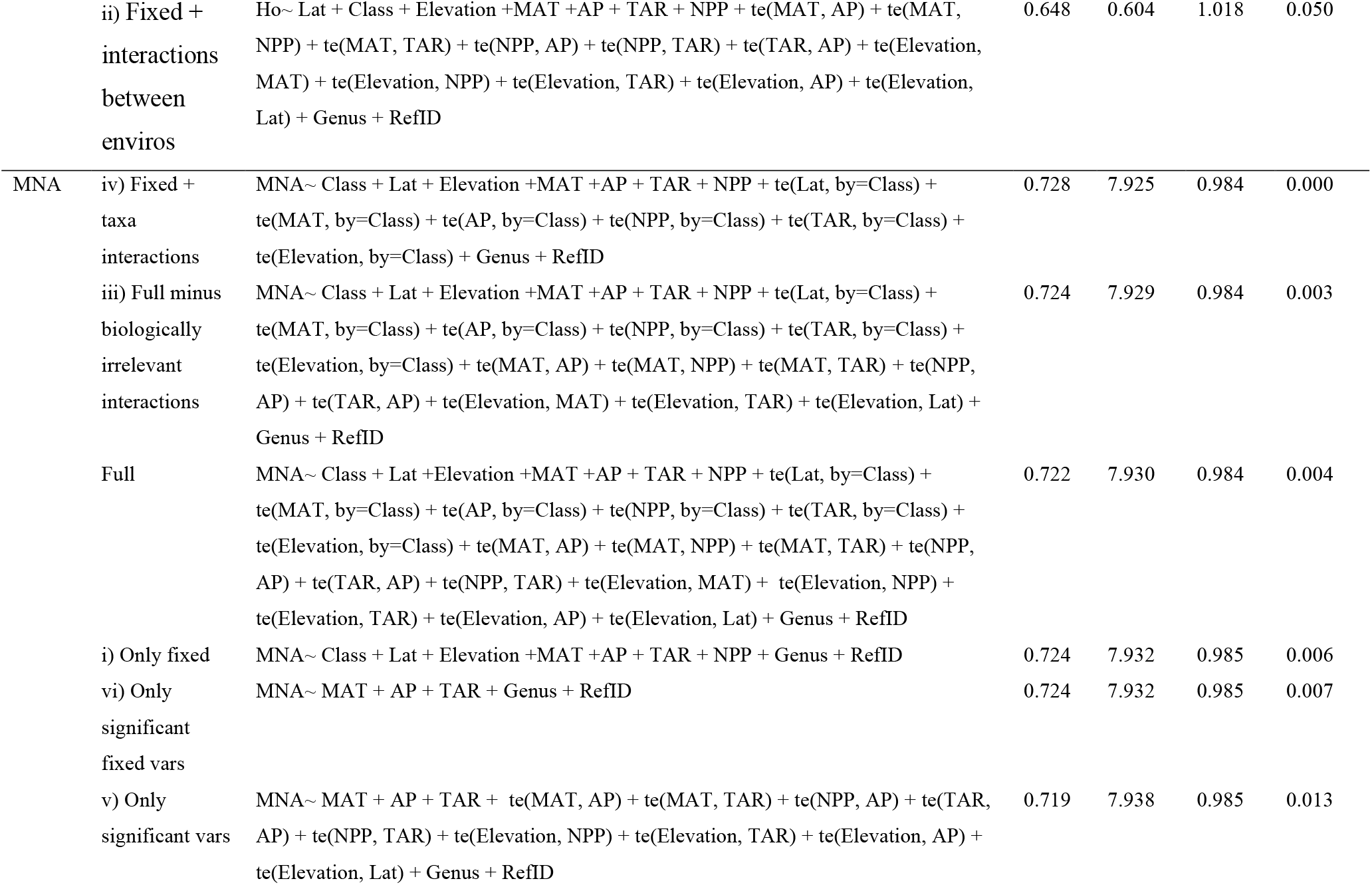
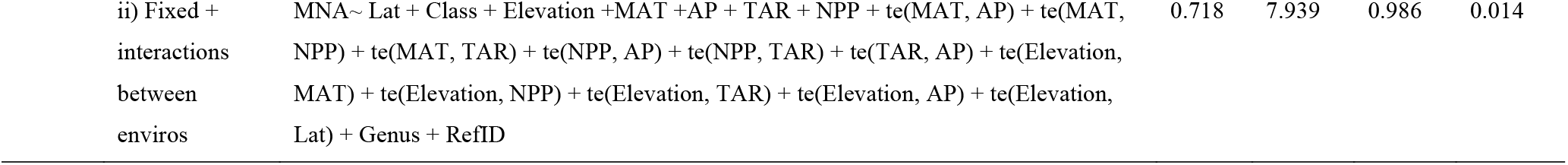
RMSE values between models. RMSE = Root Mean Square Error, PER = Prediction Error Rate. All variables except for Class were fitted with a smoothing parameter (not indicated here), interactions were fit with tensor products (te). See Table S2 for variable acronyms; letter numbering corresponds to reduced model as described in Methods. Random effects included Genus and RefID.

| PGD<br>Metric | Description | Model | R<sup>2</sup> | RMSE | PER | Delta<br>RMSE |
| --- | --- | --- | --- | --- | --- | --- |
| Ho | i) Only fixed | Ho~ Class + Lat + Elevation + MAT +AP + TAR + NPP + Genus +<br>RefID | 0.682 | 0.554 | 0.934 | 0 |
|  | iv) Fixed +<br>taxa<br>interactions | Ho~ Lat + Class + Elevation +MAT +AP + TAR + NPP + te(Lat,<br>by=Class) + te(MAT, by=Class) + te(AP, by=Class) + te(NPP, by=Class) +<br>te(TAR, by=Class) + te(Elevation, by=Class) + Genus + RefID | 0.695 | 0.569 | 0.960 | 0.016 |
|  | iii) Full minus<br>biologically<br>irrelevant<br>interactions | Ho~ Lat + Class + Elevation +MAT +AP + TAR + NPP + te(Lat,<br>by=Class) + te(MAT, by=Class) + te(AP, by=Class) + te(NPP, by=Class) +<br>te(TAR, by=Class) + te(Elevation, by=Class) + te(MAT, AP) + te(MAT,<br>NPP) + te(MAT, TAR) + te(NPP, AP) + te(TAR, AP) + te(Elevation,<br>MAT) + te(Elevation, TAR) + te(Elevation, Lat) + Genus + RefID | 0.669 | 0.588 | 0.992 | 0.034 |
|  | Full | Ho~ Lat + Class + Elevation +MAT +AP + TAR + NPP + te(Lat,<br>by=Class) + te(MAT, by=Class) + te(AP, by=Class) + te(NPP, by=Class) +<br>te(TAR, by=Class) + te(Elevation, by=Class) + te(MAT, AP) + te(MAT,<br>NPP) + te(MAT, TAR) + te(NPP, AP) + te(TAR, AP) + te(NPP, TAR) +<br>te(Elevation, MAT) + te(Elevation, NPP) + te(Elevation, TAR) +<br>te(Elevation, AP) + te(Elevation, Lat) + Genus + RefID | 0.666 | 0.598 | 1.009 | 0.044 |
|  | ii) Fixed + interactions between enviros | Ho~ Lat + Class + Elevation +MAT +AP + TAR + NPP + te(MAT, AP) + te(MAT, NPP) + te(MAT, TAR) + te(NPP, AP) + te(NPP, TAR) + te(TAR, AP) + te(Elevation, MAT) + te(Elevation, NPP) + te(Elevation, TAR) + te(Elevation, AP) + te(Elevation, Lat) + Genus + RefID | 0.648 | 0.604 | 1.018 | 0.050 |
| MNA | iv) Fixed + taxa interactions | MNA~ Class + Lat + Elevation +MAT +AP + TAR + NPP + te(Lat, by=Class) + te(MAT, by=Class) + te(AP, by=Class) + te(NPP, by=Class) + te(TAR, by=Class) + te(Elevation, by=Class) + Genus + RefID | 0.728 | 7.925 | 0.984 | 0.000 |
|  | iii) Full minus biologically irrelevant interactions | MNA~ Class + Lat + Elevation +MAT +AP + TAR + NPP + te(Lat, by=Class) + te(MAT, by=Class) + te(AP, by=Class) + te(NPP, by=Class) + te(TAR, by=Class) + te(Elevation, by=Class) + te(MAT, AP) + te(MAT, NPP) + te(MAT, TAR) + te(NPP, AP) + te(TAR, AP) + te(Elevation, MAT) + te(Elevation, TAR) + te(Elevation, Lat) + Genus + RefID | 0.724 | 7.929 | 0.984 | 0.003 |
|  | Full | MNA~ Class + Lat +Elevation +MAT +AP + TAR + NPP + te(Lat, by=Class) + te(MAT, by=Class) + te(AP, by=Class) + te(NPP, by=Class) + te(TAR, by=Class) + te(Elevation, by=Class) + te(MAT, AP) + te(MAT, NPP) + te(MAT, TAR) + te(NPP, AP) + te(TAR, AP) + te(NPP, TAR) + te(Elevation, MAT) + te(Elevation, NPP) + te(Elevation, TAR) + te(Elevation, AP) + te(Elevation, Lat) + Genus + RefID | 0.722 | 7.930 | 0.984 | 0.004 |
|  | i) Only fixed | MNA~ Class + Lat + Elevation +MAT +AP + TAR + NPP + Genus + RefID | 0.724 | 7.932 | 0.985 | 0.006 |
|  | vi) Only significant fixed vars | MNA~ MAT + AP + TAR + Genus + RefID | 0.724 | 7.932 | 0.985 | 0.007 |
|  | v) Only significant vars | MNA~ MAT + AP + TAR + te(MAT, AP) + te(MAT, TAR) + te(NPP, AP) + te(TAR, AP) + te(NPP, TAR) + te(Elevation, NPP) + te(Elevation, TAR) + te(Elevation, AP) + te(Elevation, Lat) + Genus + RefID | 0.719 | 7.938 | 0.985 | 0.013 |

|  |  |  |  |  |  |
| --- | --- | --- | --- | --- | --- |
| ii) Fixed +<br>interactions<br>between<br>enviros | MNA~ Lat + Class + Elevation +MAT +AP + TAR + NPP + te(MAT, AP) + te(MAT,<br>NPP) + te(MAT, TAR) + te(NPP, AP) + te(NPP, TAR) + te(TAR, AP) + te(Elevation,<br>MAT) + te(Elevation, NPP) + te(Elevation, TAR) + te(Elevation, AP) + te(Elevation,<br>Lat) + Genus + RefID | 0.718 | 7.939 | 0.986 | 0.014 |

**Table S4.** Summary of mean environmental and population genetic variables for each of the vertebrate classes assessed across the Americas (before separating into Ho and MNA datasets). Ho = observed heterozygosity, MNA = mean number of alleles, Lat = degrees latitude, MAT = mean annual temperature (°C), AP = annual precipitation (mm/year), TAR = total annual range (°C), Elevation (m), NPP = net primary productivity (units of elemental carbon x10e^-11^). Values in parentheses represent the standard deviation.

| Variable | Amphibia | Anadromous | Aves | Freshwater | Mammalia | Reptilia |
| --- | --- | --- | --- | --- | --- | --- |
| # populations | 1117 | 1315 | 608 | 2700 | 1943 | 1349 |
| MNA | 7.49<br>(5.56) | 14.97<br>(7.01) | 7.65<br>(6.16) | 7.11<br>(4.01) | 6.09<br>(3.93) | 6.69<br>(3.63) |
| Ho | 0.57<br>(0.17) | 0.70<br>(0.15) | 0.59<br>(0.17) | 0.57<br>(0.17) | 0.60<br>(0.14) | 0.59<br>(0.15) |
| Lat | 32.72<br>(18.27) | 50.55<br>(5.95) | 25.88<br>(26.03) | 37.44<br>(20.42) | 34.19<br>(27.15) | 27.54<br>(16.37) |
| MAT | 12.51<br>(7.25) | 7.35<br>(4.19) | 14.25<br>(8.86) | 10.46<br>(8.24) | 9.43<br>(9.10) | 17.97<br>(6.56) |
| AP | 1258.83<br>(893.63) | 1122.44<br>(743.53) | 1230.72<br>(866.84) | 988.72<br>(593.08) | 917.45<br>(739.85) | 1113.82<br>(651.43) |
| TAR | 26.87<br>(10.64) | 26.93<br>(6.91) | 22.20<br>(12.76) | 29.89<br>(9.78) | 30.08<br>(10.17) | 23.19<br>(11.24) |
| Elevation | 805.93<br>(890.74) | 222.98<br>(338.72) | 519.63<br>(742.59) | 472.06<br>(605.76) | 736.71<br>(869.81) | 278.28<br>(441.53) |
| NPP | 3.78E+11<br>(2.16E+11) | 2.56E+11<br>(1.46E+11) | 3.25E+11<br>(2.64E+11) | 3.42E+11<br>(2.39E+11) | 2.79E+11<br>(2.16E+11) | 3.45E+11<br>(2.46E+11) |

**Table S5.**
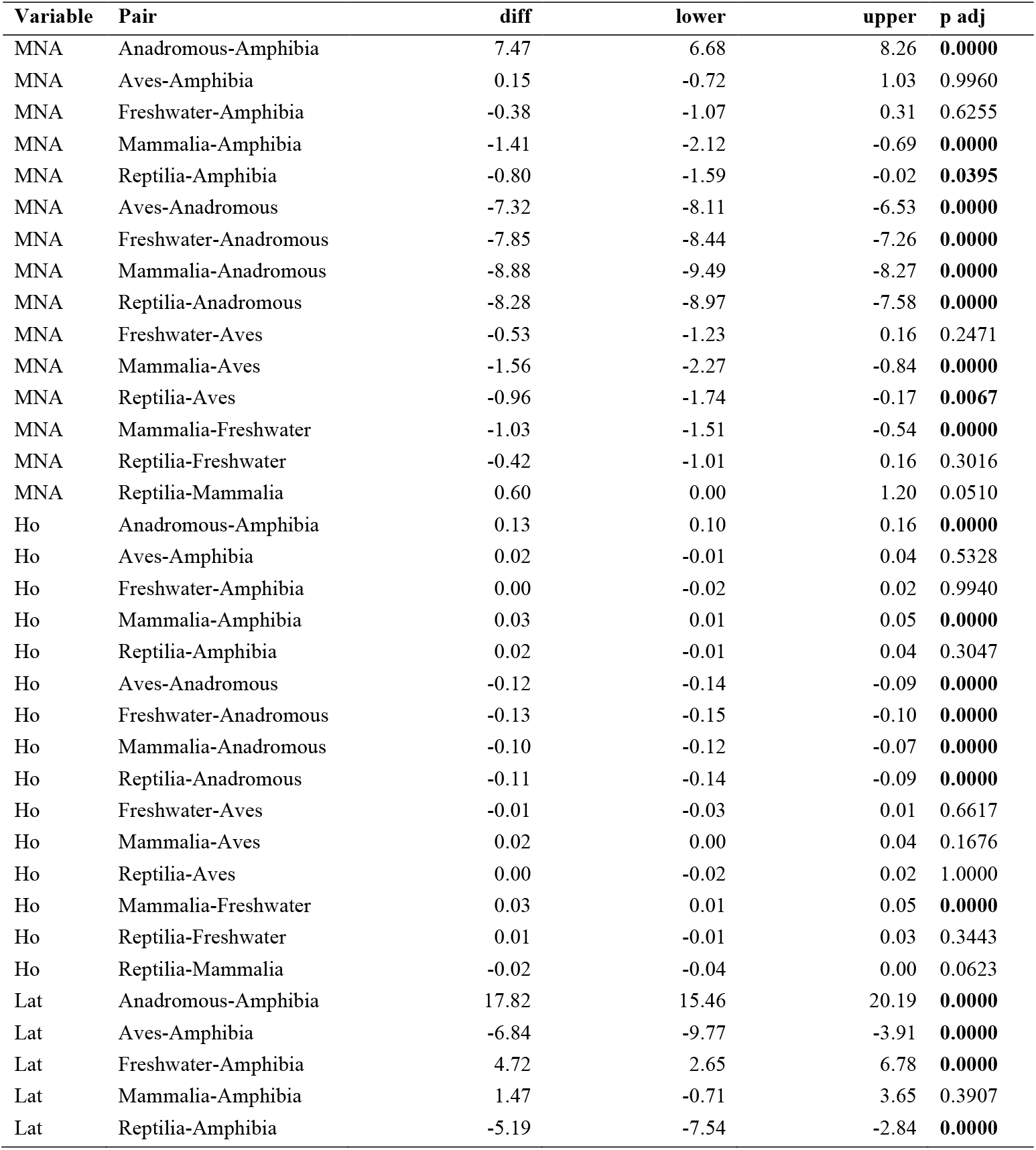
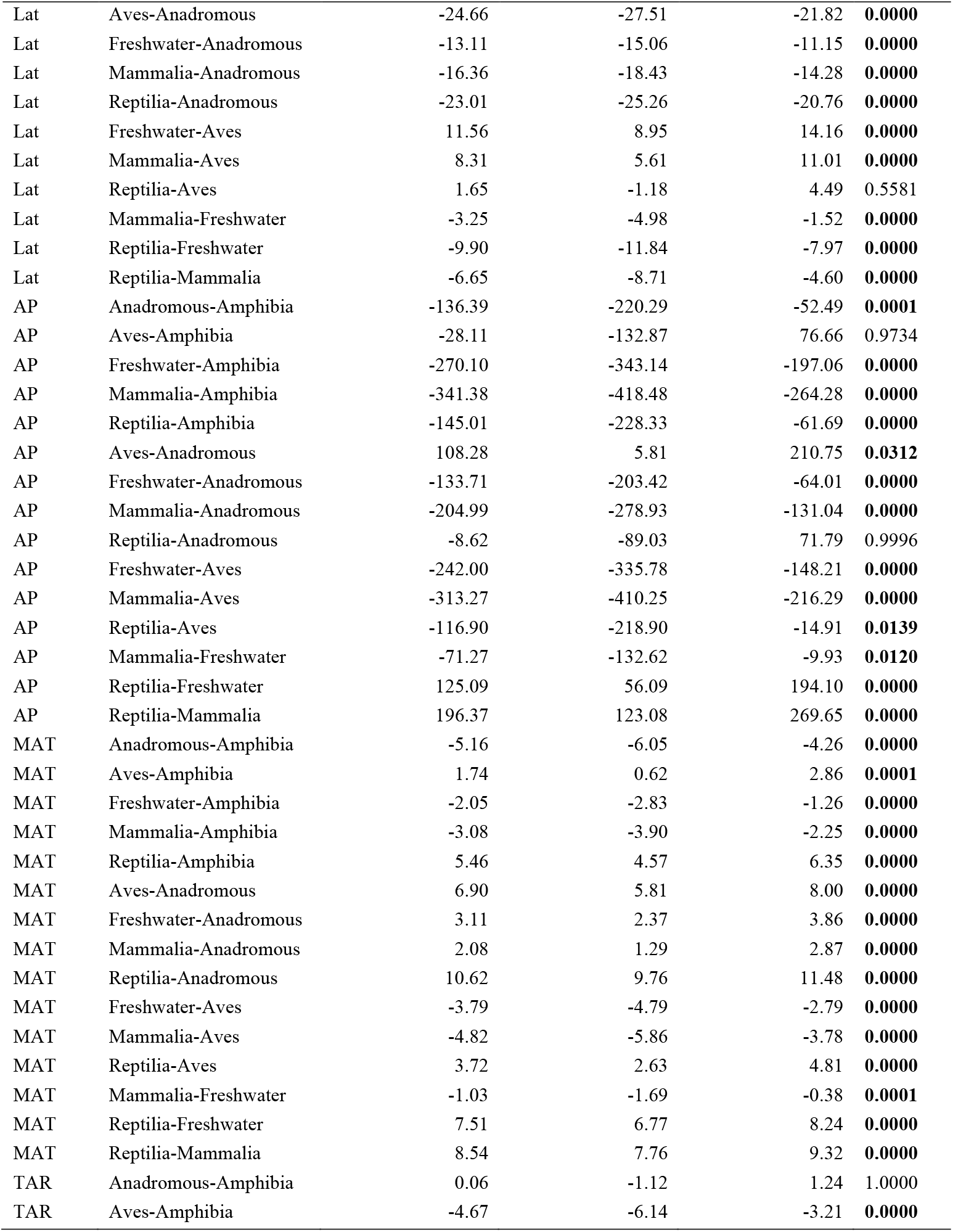
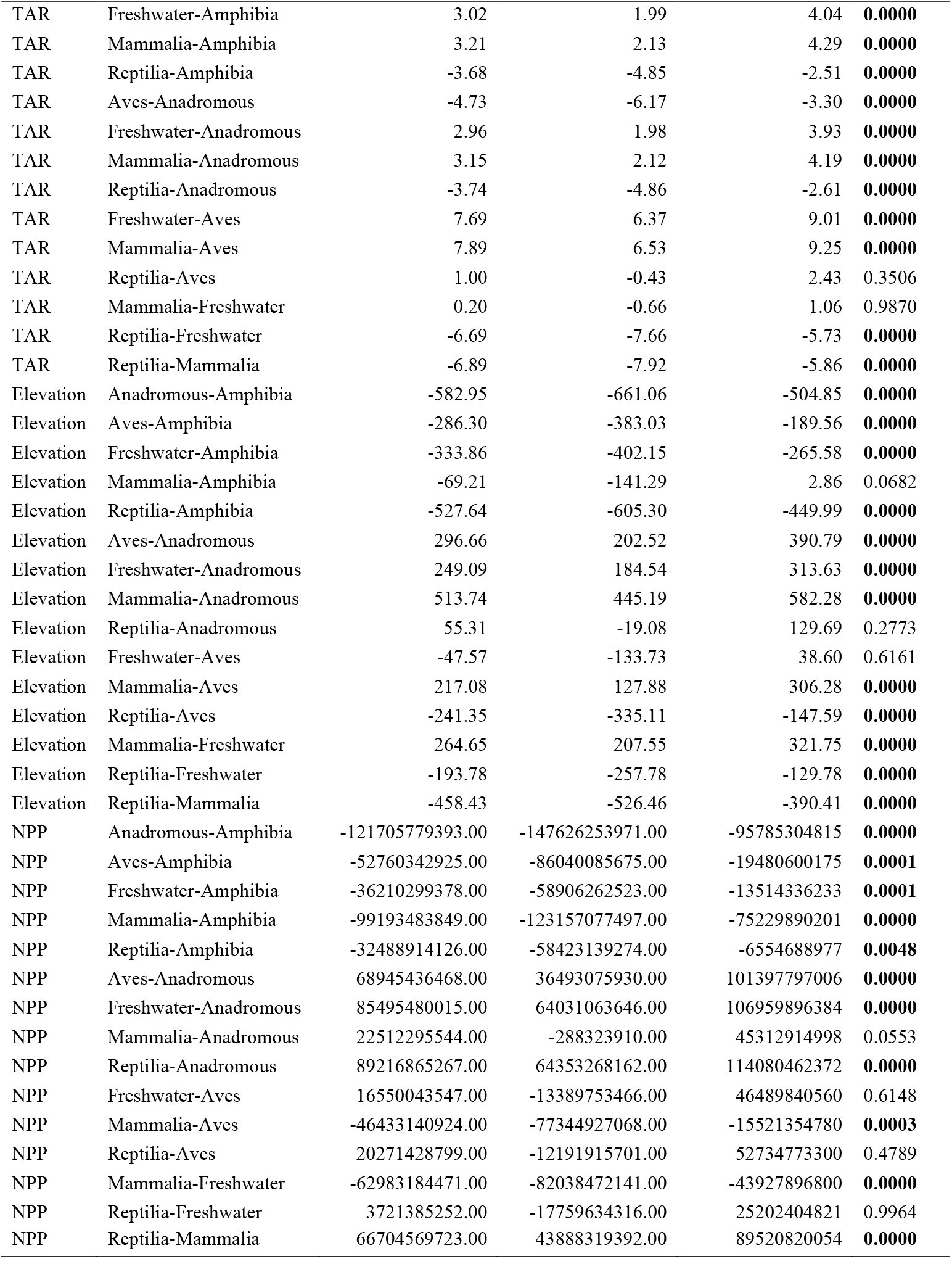
Pairwise differences among taxa for each environmental variable. Ho = observed heterozygosity, MNA = mean number of alleles, Lat = degrees latitude, MAT = mean annual temperature (°C), AP = annual precipitation (mm/year), TAR = total annual range (°C), Elevation (m), NPP = net primary productivity (units of elemental carbon x10e^-11^). Bold indicates statistical significance p<0.05.

**Figure S1.**
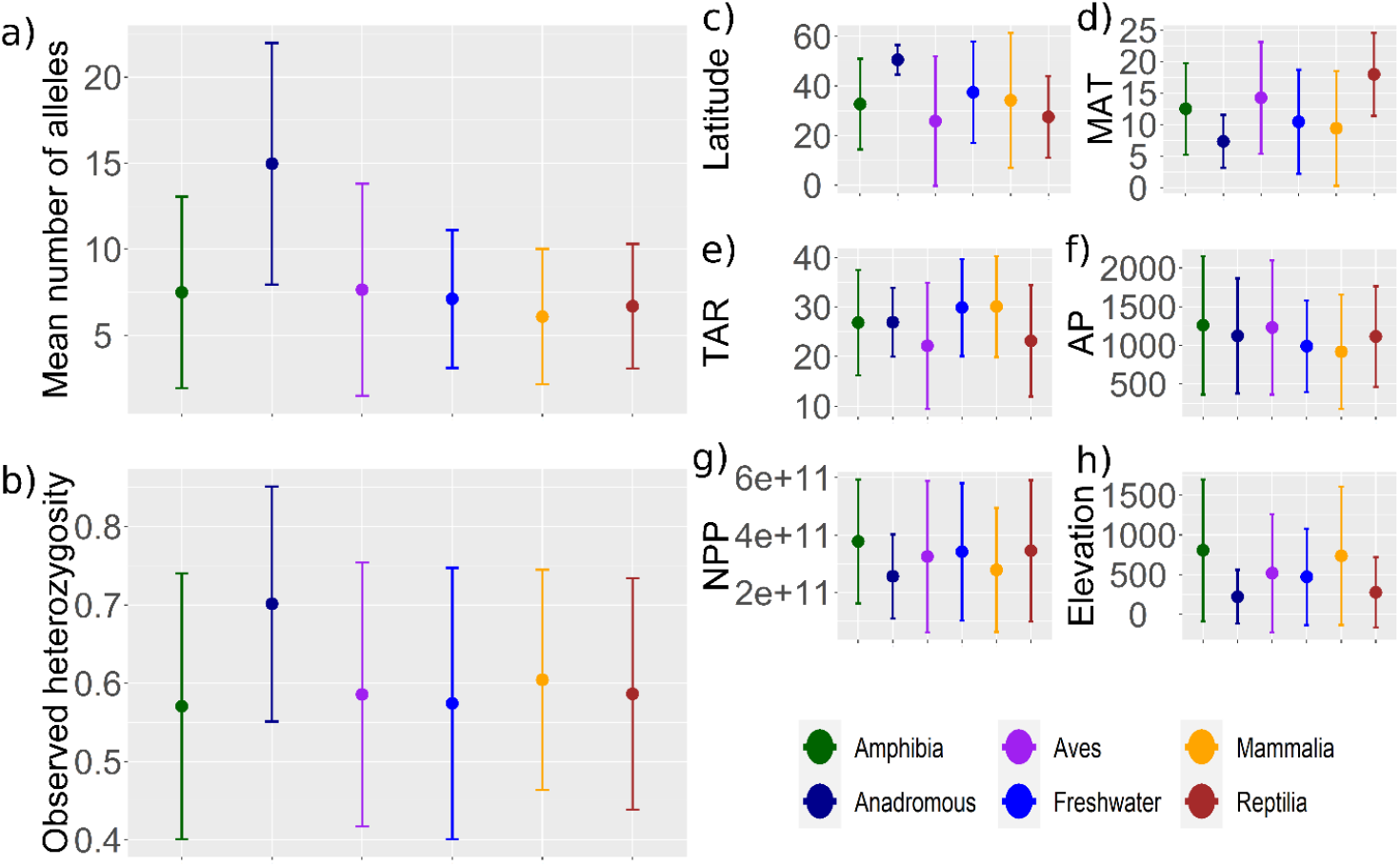
Means and standard deviations for each taxonomic Class for a) mean number of alleles (MNA), b) observed heterozygosity (Ho), c) degrees latitude, d) MAT = mean annual temperature (°C, MAT), e) TAR = temperature annual range (°C, TAR), f) AP = annual precipitation (mm/year, AP), g) NPP = net primary productivity (units elemental carbon, NPP), h) elevation (m).

**Figure S2.**
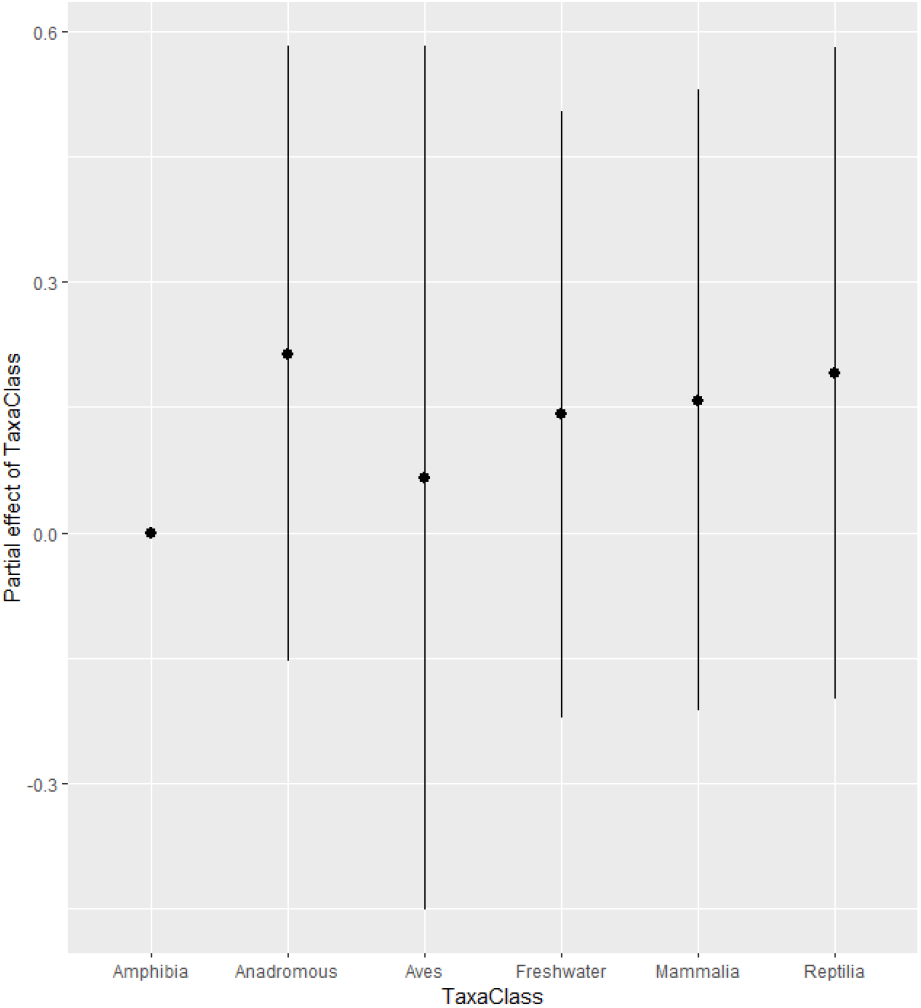
The effect of vertebrate Class for the observed heterozygosity model. Indicating no significant differences between Classes.

**Figure S3.**
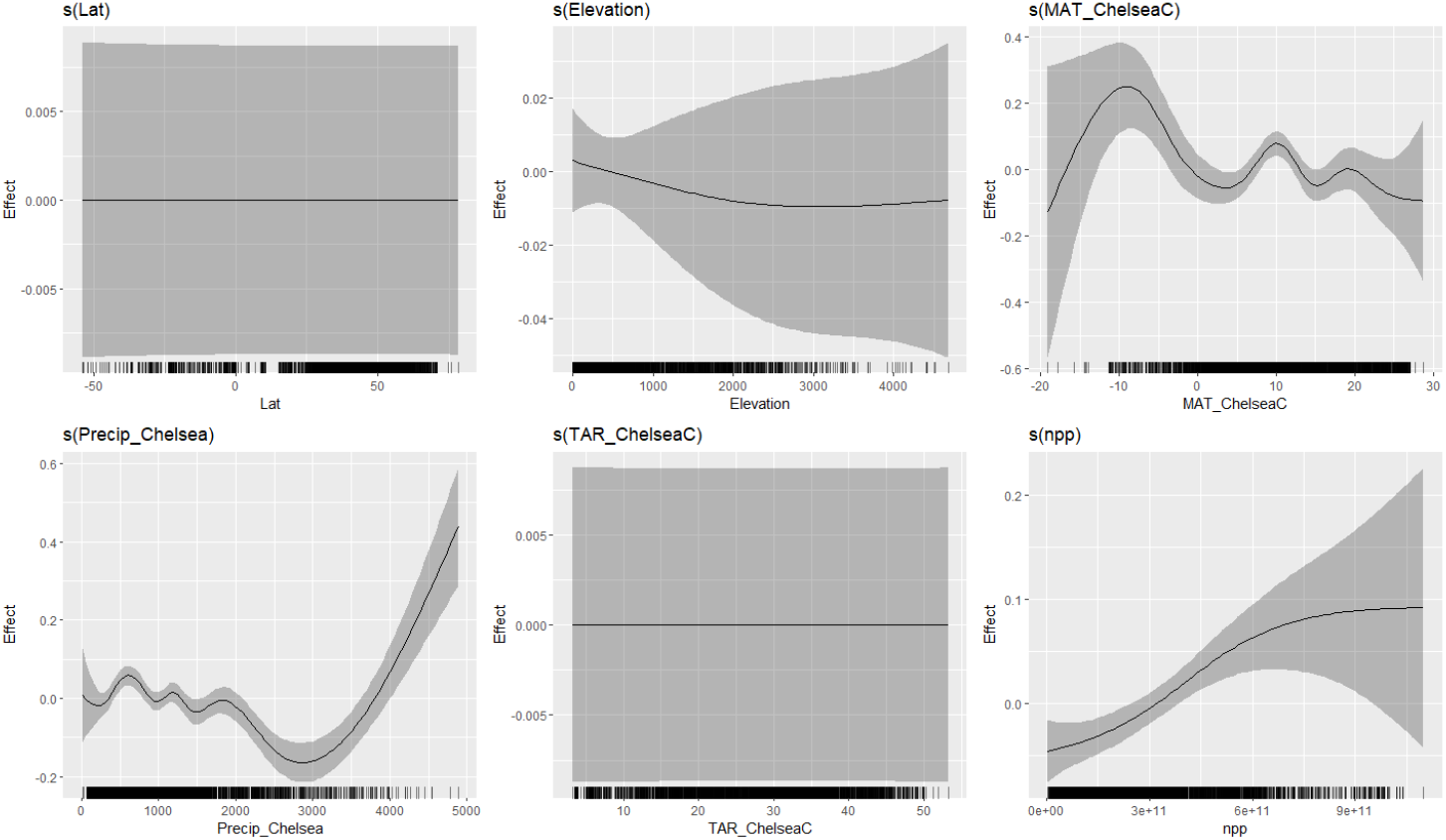
Global smoothers for the MNA model. Note the y axis does not show MNA but rather the effect of the variable on MNA, where values above or below zero represent positive or negative changes about the mean (zero), respectively.

**Figure S4.**
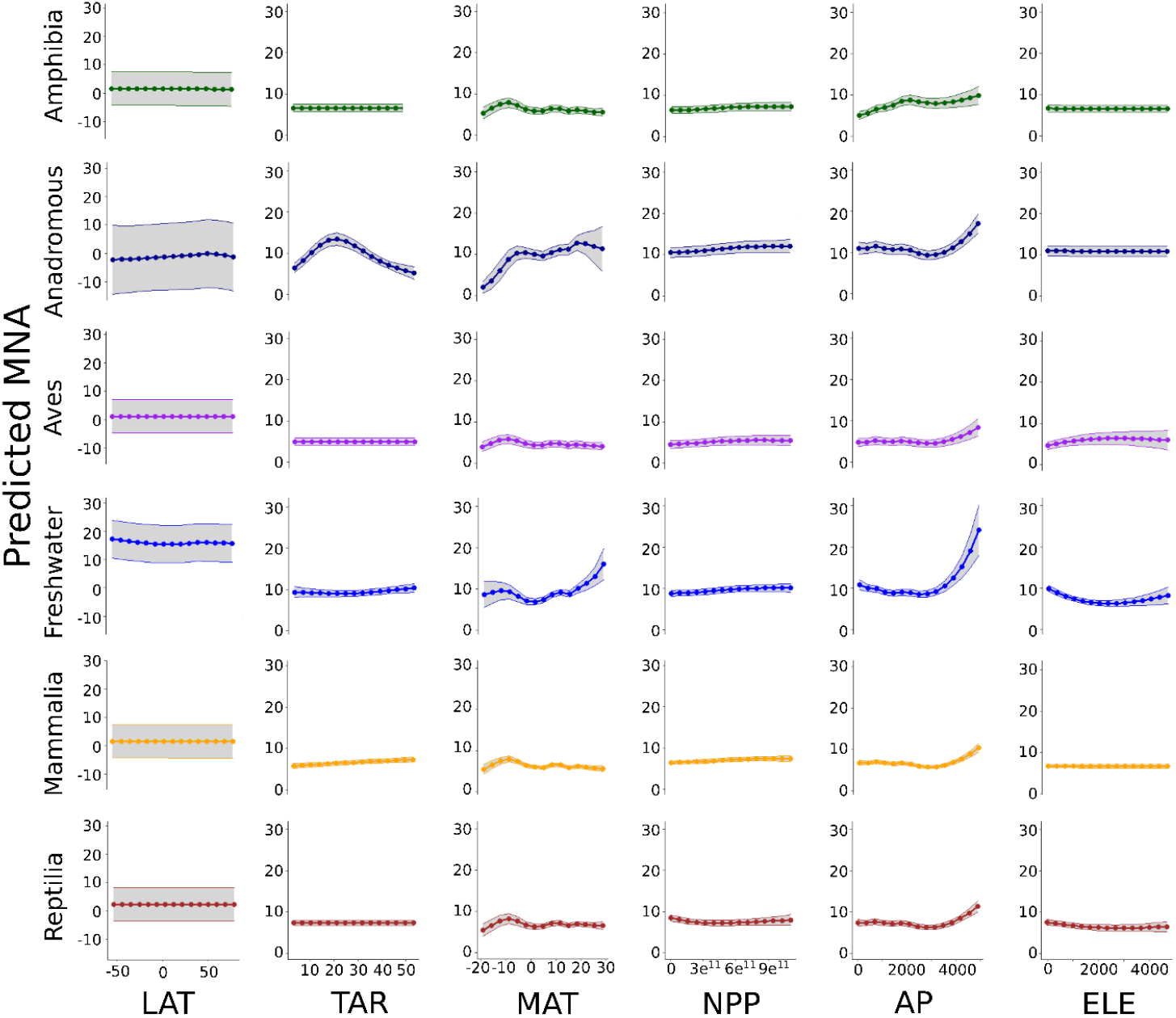
The predicted effect of latitude and environmental variables on mean number of alleles (MNA) for all vertebrate Classes across the Americas. Predictors were fitted by cubic smoothers and include, in columns from left to right: degrees latitude; TAR= total annual temperature range (°C); MAT = mean annual temperature (°C); NPP = net primary productivity (units of elemental carbon x10e^-11^; AP = annual precipitation (mm/year); ELE = Elevation (m). Colours represent the five different vertebrate Classes and lighter bands represent standard error.

**Figure S5.**
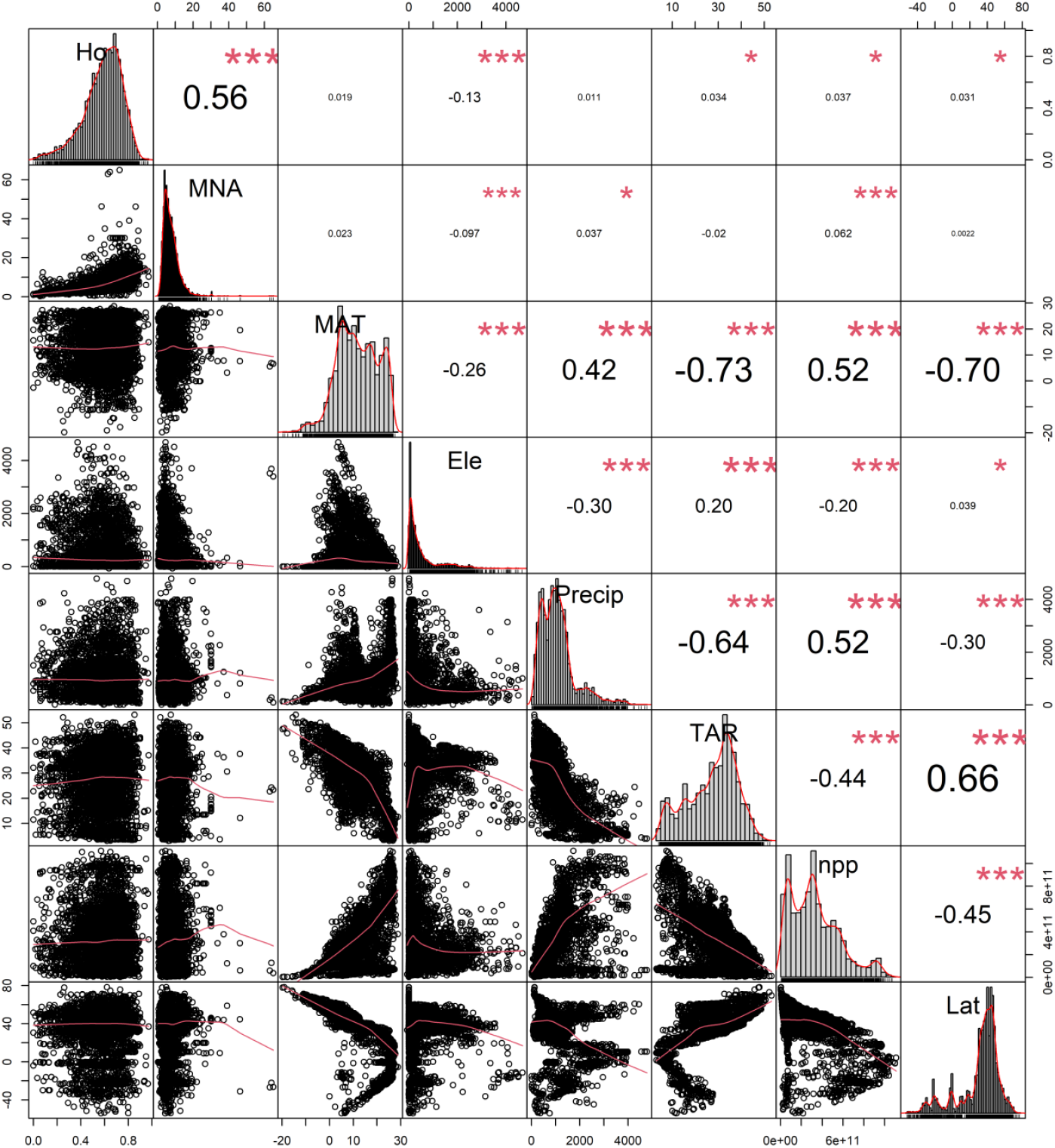
Pairwise correlation between population genetic diversity (Ho=observed heterozygosity, MNA=mean number of alleles) and environmental variables. MAT= mean annual temperature (°C), Ele = elevation (m), Precip = annual precipitation (mm/year), TAR= temperature annual range (°C), npp= net primary productivity (units elemental carbon), Lat=degrees latitude.

