## Supplemental File 2 for "Macrogenetics reveals multifaceted influences of environmental variation on vertebrate population genetic diversity across the Americas"

### Supplementary File 2 for

Multifaceted environmental and latitudinal influences on vertebrate population genetic diversity across the Americas

**Supplementary File 2: GAMM plot output for MNA**

MNA ~ s(Lat, bs = "cs") + s(Elevation, bs = "cs") +

s(MAT_ChelseaC, bs = "cs") + s(Precip_Chelsea, bs = "cs") +

s(TAR_ChelseaC, bs = "cs") + s(npp, bs = "cs") +

te(Lat, by = TaxaClass, m = 1, bs = "cs") + te(MAT_ChelseaC,

by = TaxaClass, m = 1, bs = "cs") + te(Precip_Chelsea,

by = TaxaClass, m = 1, bs = "cs") + te(npp, by = TaxaClass,

m = 1, bs = "cs") + te(TAR_ChelseaC, by = TaxaClass,

m = 1, bs = "cs") + te(Elevation, by = TaxaClass, m = 1,

bs = "cs") + te(MAT_ChelseaC, Precip_Chelsea, m = 1,

bs = "cs") + te(MAT_ChelseaC, npp, m = 1, bs = "cs") +

te(MAT_ChelseaC, TAR_ChelseaC, m = 1, bs = "cs") +

te(Elevation, MAT_ChelseaC, m = 1, bs = "cs") + te(Elevation,

npp, m = 1, bs = "cs") + te(Elevation, TAR_ChelseaC,

m = 1, bs = "cs") + te(Elevation, Precip_Chelsea, m = 1,

bs = "cs") + te(Elevation, Lat, m = 1, bs = "cs") +

te(npp, Precip_Chelsea, m = 1, bs = "cs") + te(npp,

TAR_ChelseaC, m = 1, bs = "cs") + te(TAR_ChelseaC,

Precip_Chelsea, m = 1, bs = "cs") + s(Genus, bs = "re") +

s(RefID, bs = "re") + TaxaClass

**Variables on their own**

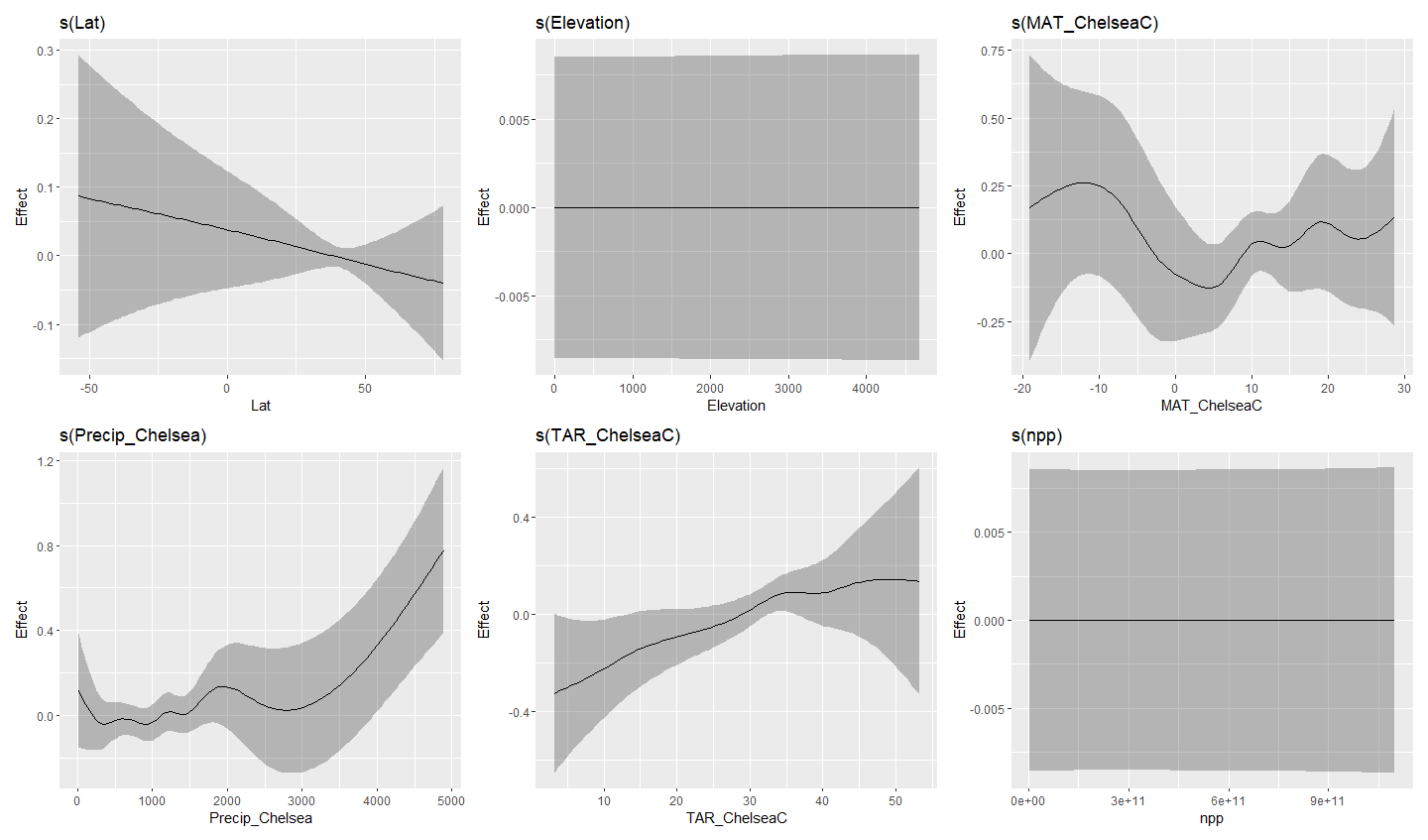

**Class:Latitude**

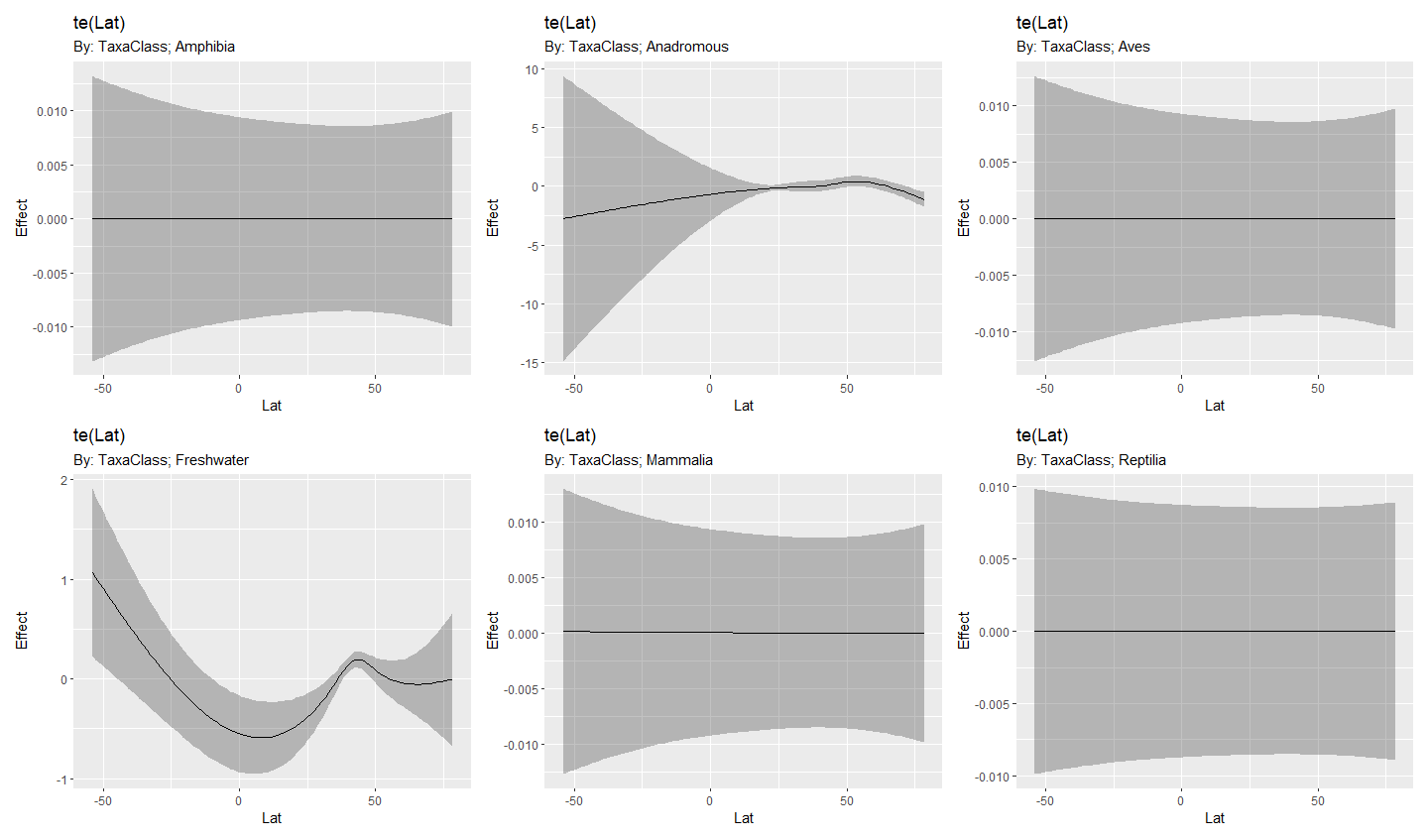

**Class:MAT**

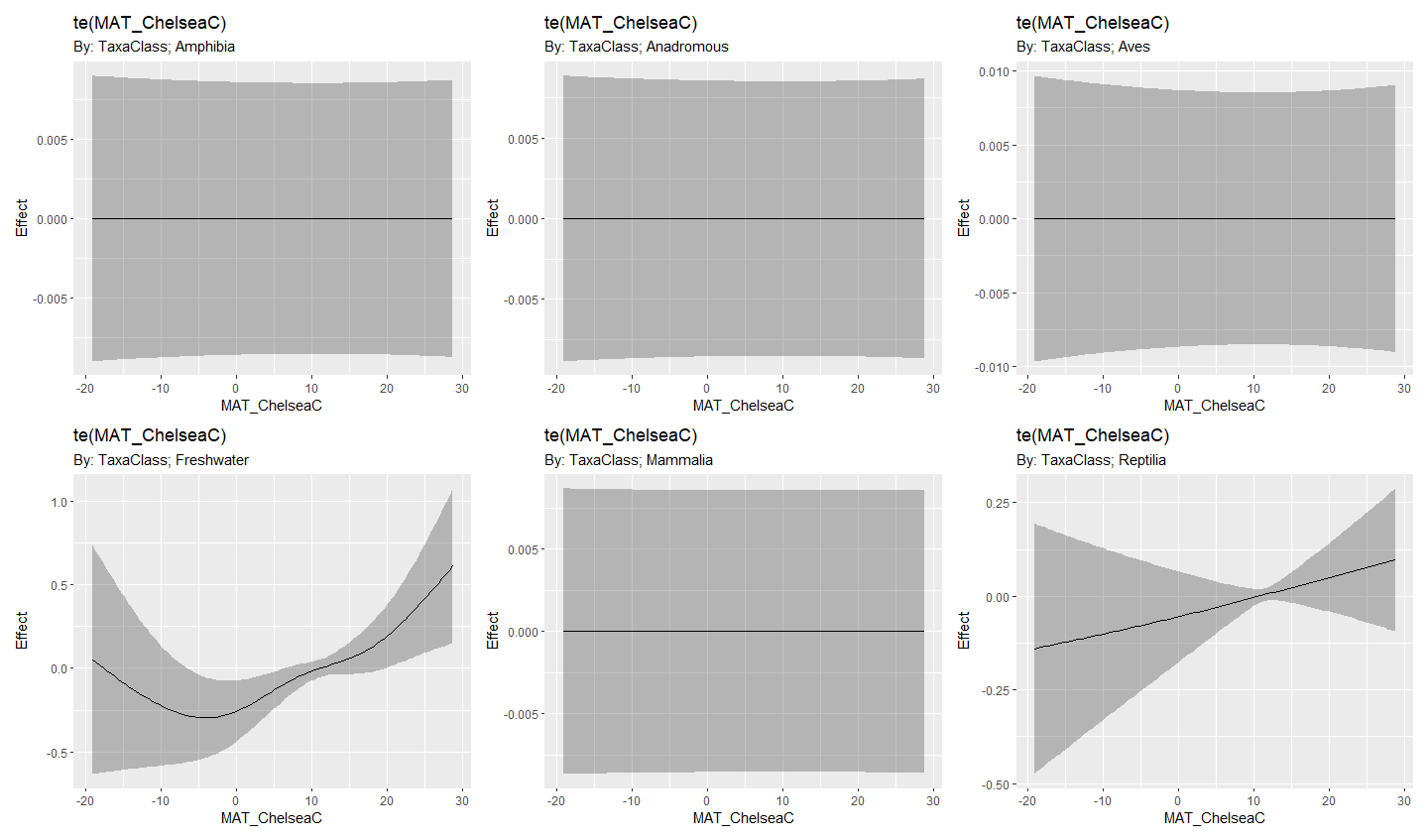

**Class:Precip**

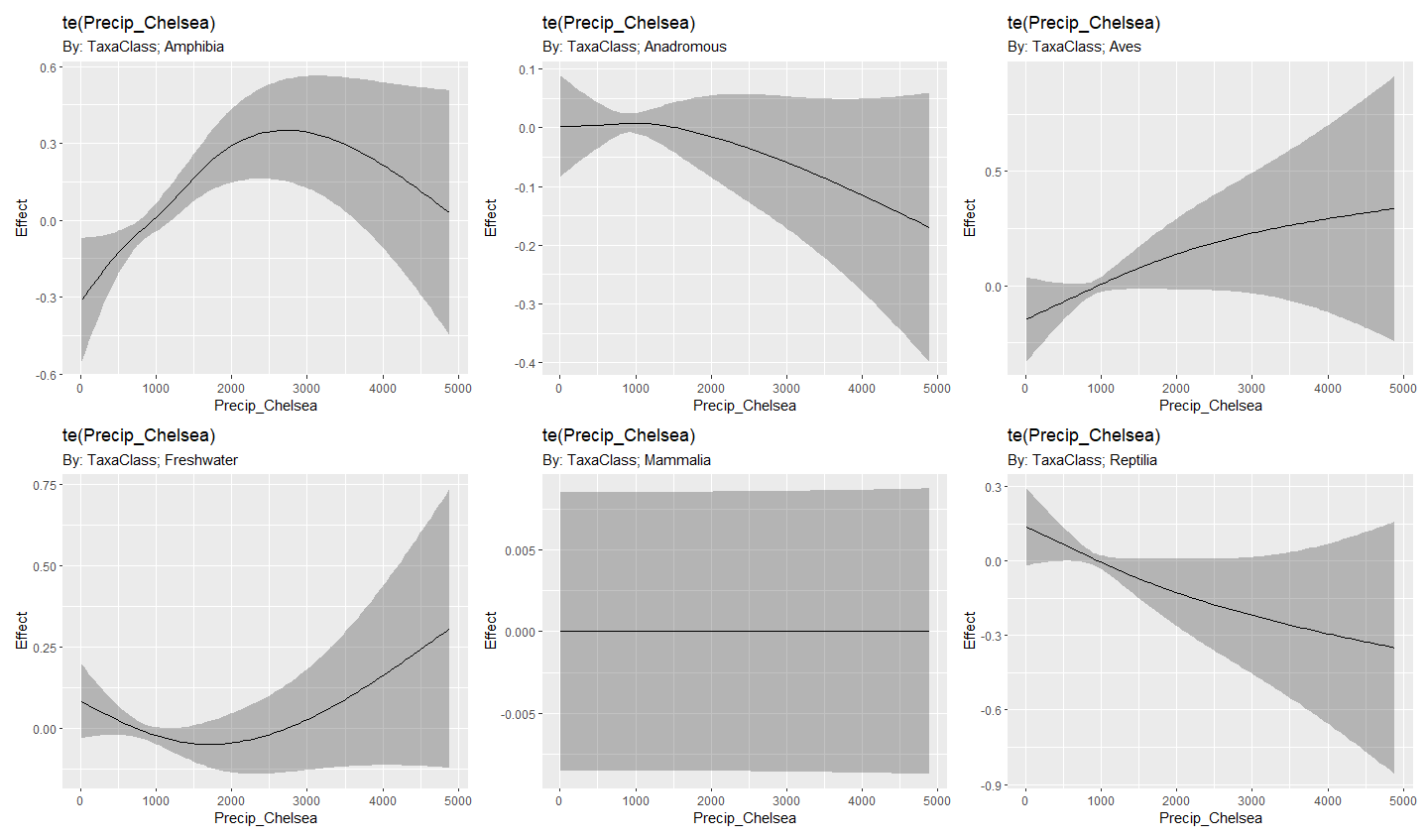

**Class:NPP**

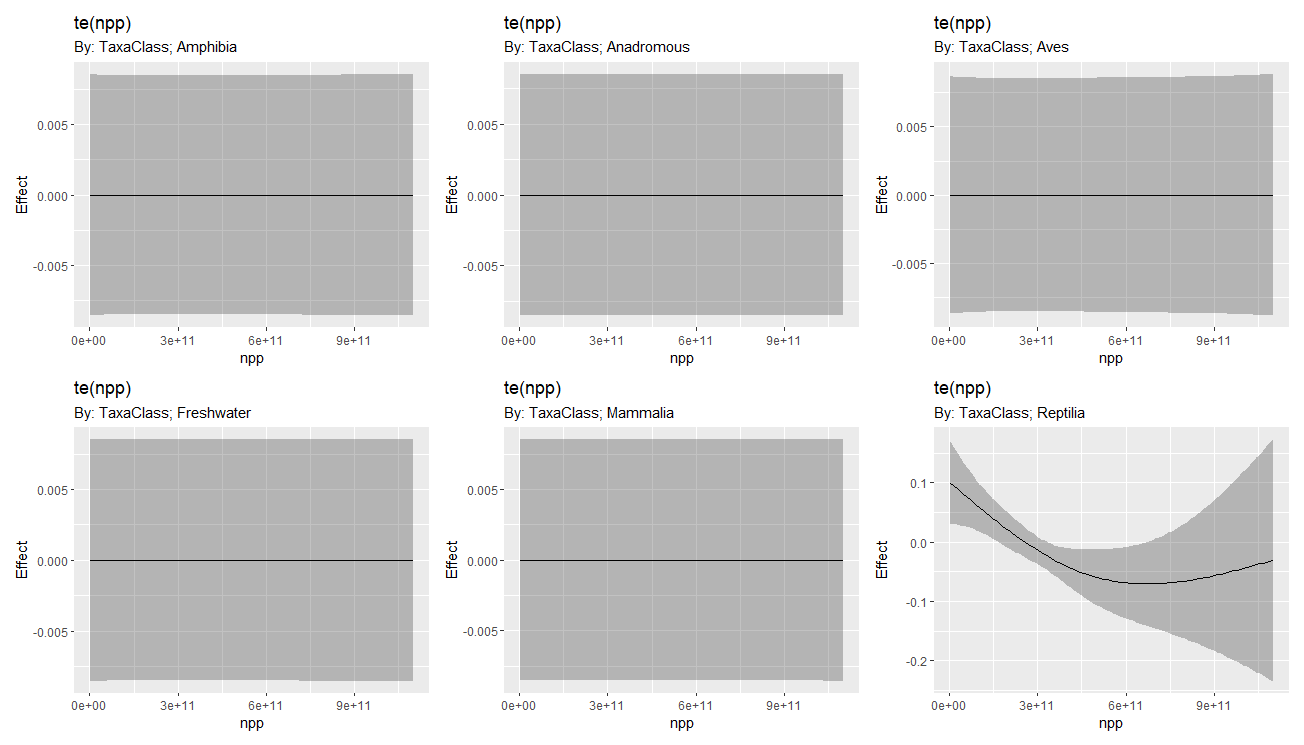

**Class:TAR**

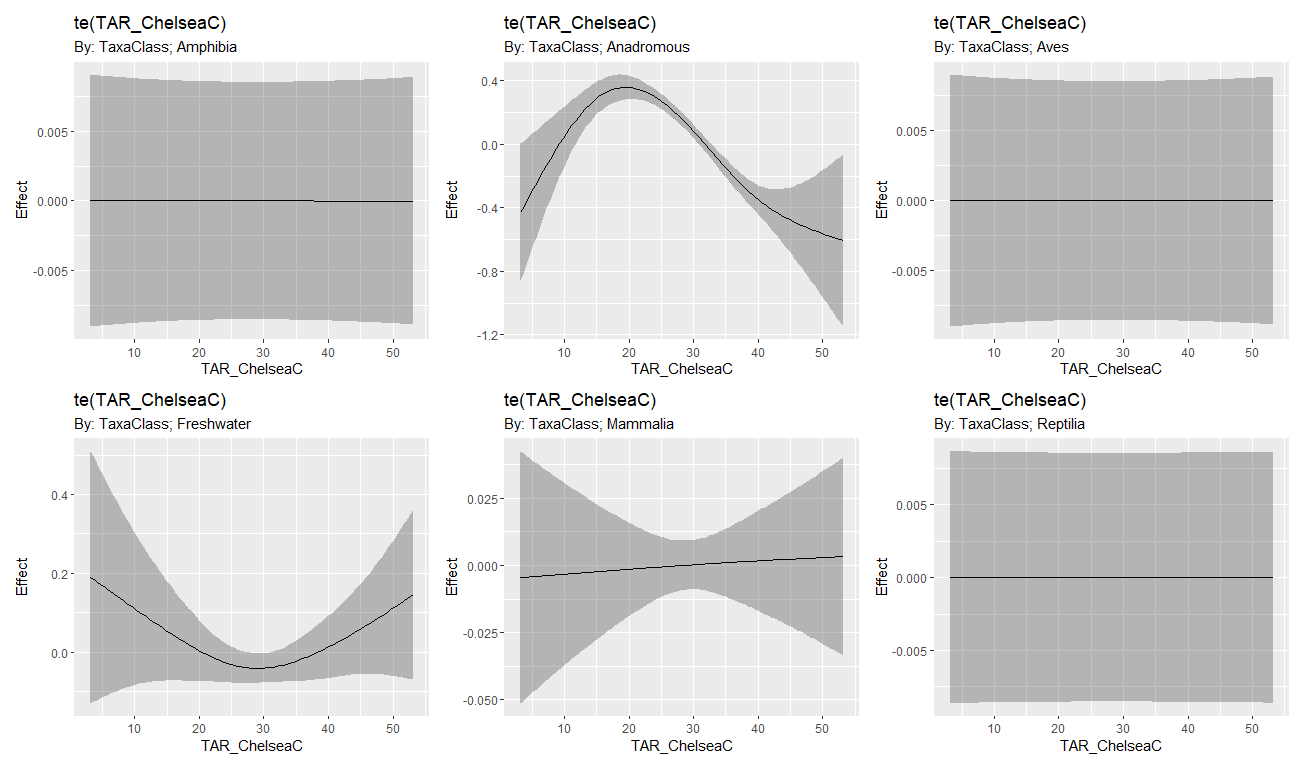

**Class:Elevation**

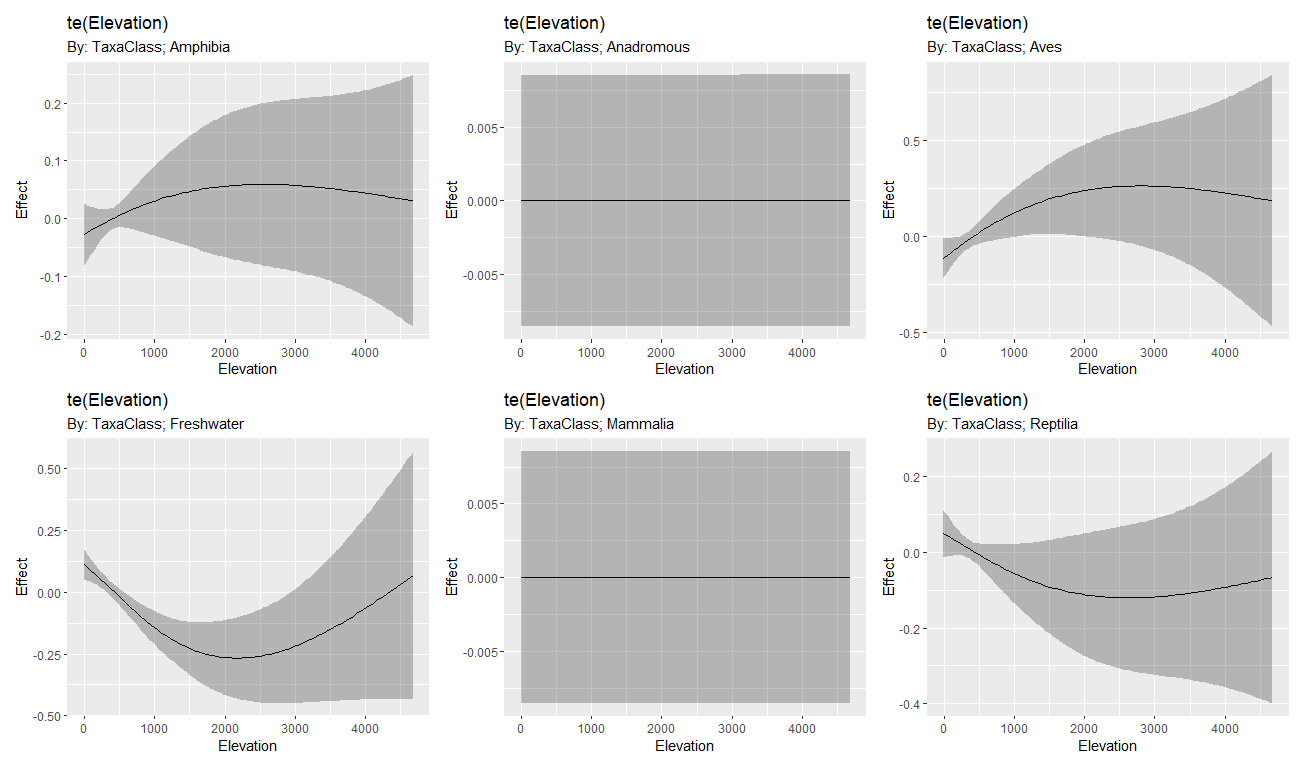

**MAT with other environmental variables**

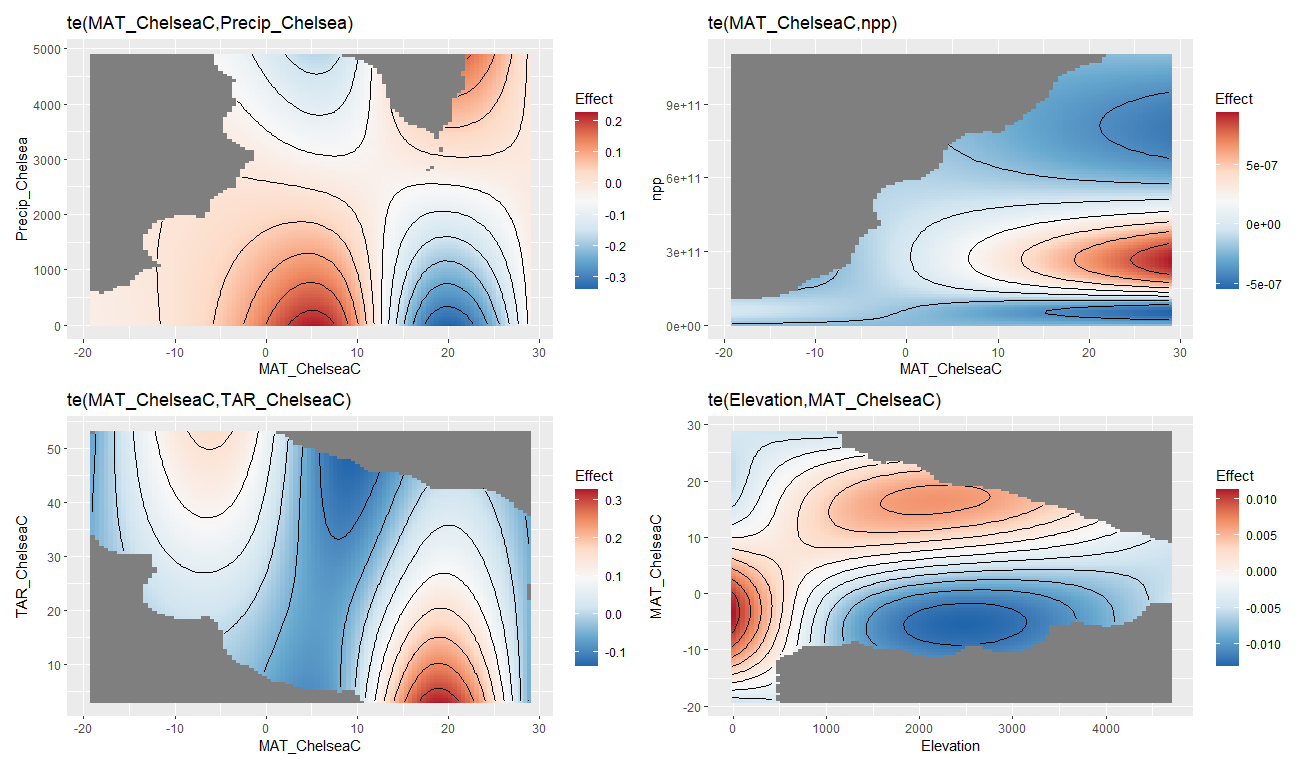

**Elevation with other environmental variables**

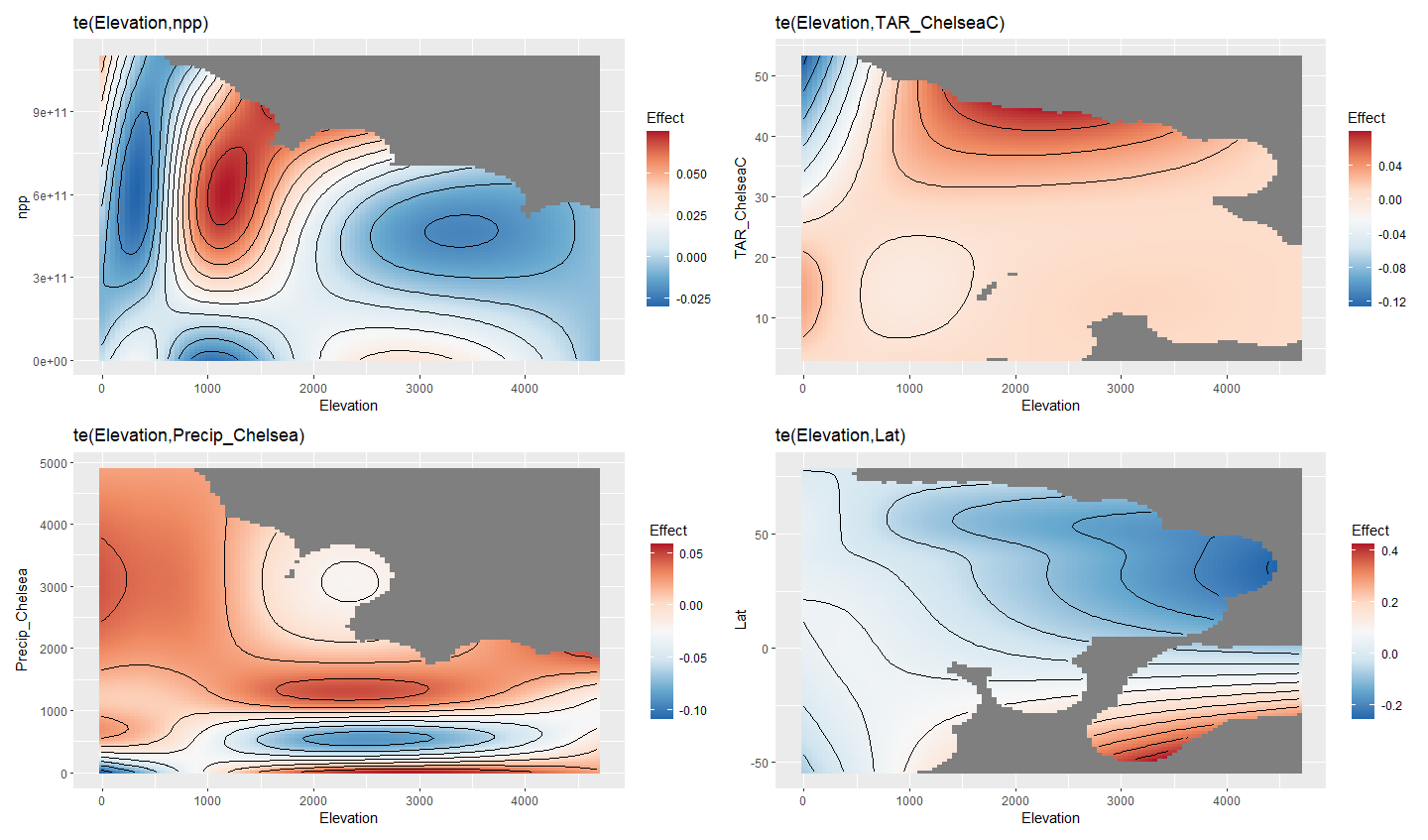

**Other interactions + randoms**

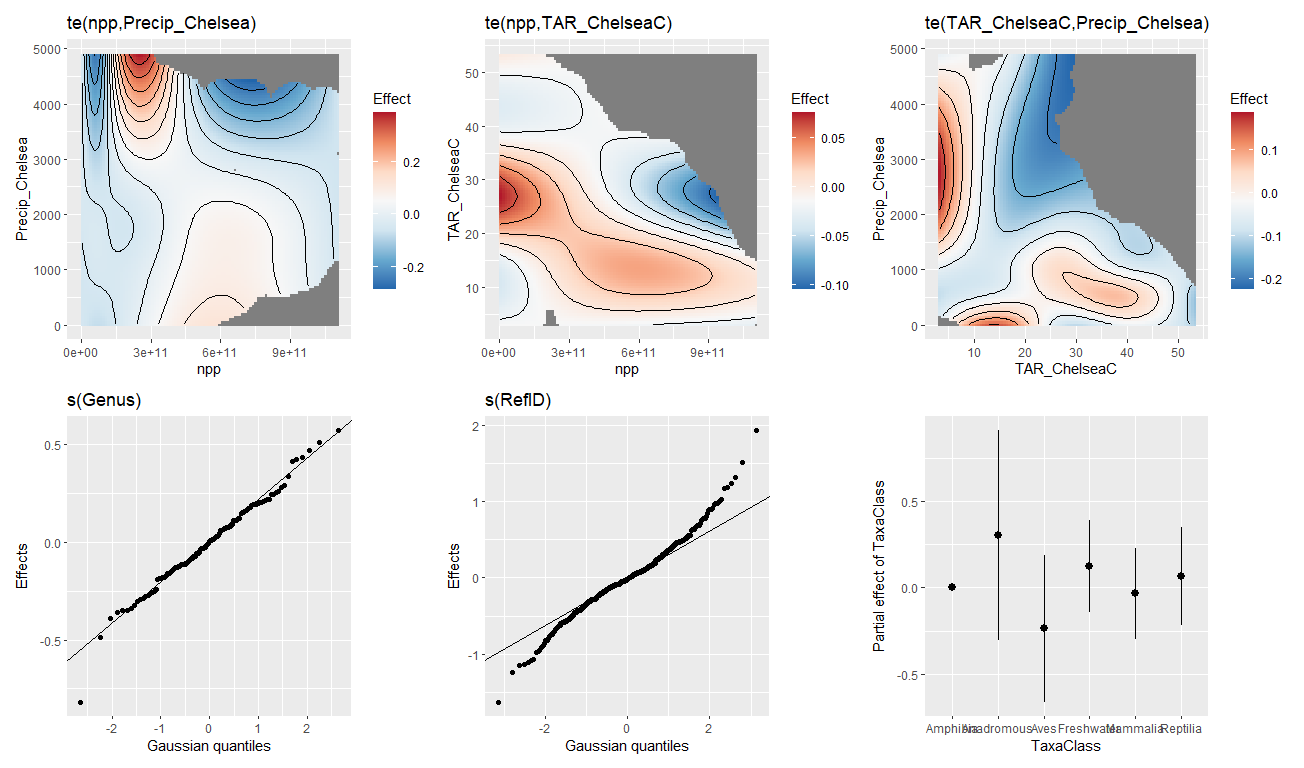
